# HAP40 is a conserved central regulator of Huntingtin and a specific modulator of mutant Huntingtin toxicity

**DOI:** 10.1101/2020.05.27.119552

**Authors:** Shiyu Xu, Gang Li, Xin Ye, Dongsheng Chen, Zhihua Chen, Zhen Xu, Lili Ye, Erin F. Stimming, Deanna Marchionini, Sheng Zhang

## Abstract

Perturbation of Huntingtin (HTT)’s physiological function is one postulated pathogenic factor in Huntington’s disease (HD). However, little is known how HTT is regulated *in vivo*. In a proteomic study, we isolated a novel ∼40kDa protein as a strong binding partner of *Drosophila* HTT and demonstrated it was the functional ortholog of HAP40, an HTT associated protein shown recently to modulate HTT’s conformation but with unclear physiological and pathologic roles. We showed that in both flies and human cells, HAP40 maintained conserved physical and functional interactions with HTT, loss of HAP40 resulted in similar phenotypes as HTT knockout, including animal viability and autophagy, and more strikingly, HAP40 depletion significantly reduced the levels of endogenous HTT, while HAP40 was mostly degraded via the proteasome in the absence of HTT. Interestingly, polyglutamine expansion in HTT did not affect its affinity for HAP40. However, HAP40 modulated HD pathogenesis in *Drosophila* model by regulating the overall protein levels and the toxicity of full-length mutant HTT. Together, our study uncovers a conserved mechanism governing the stability and *in vivo* functions of HTT, and demonstrates that HAP40 is a central and positive regulator of HTT, a potential modulator of HD pathogenesis and a promising candidate for “HTT-lowering” strategy against HD.

## Introduction

Huntington’s disease (HD) is a neurodegenerative disorder caused by an abnormal expansion of a glutamine tract (polyQ) near the N-terminus of Huntingtin (HTT) protein (The Huntington’s Disease Collaborative Research Group. 1993). Despite its simple genetic cause, HD has a rather complex etiology that remains to be elucidated (Bates et al., 2015; Ross et al., 2014). In particular, although HTT gene is broadly expressed throughout the brain, HD preferentially destroys medium spiny neurons in the striatum and pyramidal neurons in the cortex (Gutekunst et al., 1995; Li et al., 1993; Sharp et al., 1995; Strong et al., 1993; Trottier et al., 1995). This and other evidence have led to the hypothesis that longer polyQ track not only confers a dominant toxicity, but also perturbs HTT’s normal physiology functions, and these combinatorial attacks together lead to selective neuronal loss over times (Cattaneo et al., 2005; Saudou and Humbert, 2016). Because of this, there is growing interest in the “HTT-lowering” strategy, which targets HTT itself to eliminate mutant HTT specifically or to lower the levels of total HTT, as a promising therapeutic strategy against HD (Tabrizi et al., 2019). Thus, a clear understanding on how endogenous HTT is regulated *in vivo*, including its protein stability and its normal cellular functions, are critical both for elucidating HD etiology and for identifying effective drug targets.

HTT is a widely expressed, large ∼350 kDa protein (Gutekunst et al., 1995; Sharp et al., 1995; Trottier et al., 1995). In mouse, it is essential for early embryogenesis and for neuronal survival in postnatal brain (Duyao et al., 1995; Metzler et al., 2000; Nasir et al., 1995; Reiner et al., 2003; Zeitlin et al., 1995). Structurally, HTT is composed primarily of HEAT repeats, an anti-parallel double-helix motif implicated in protein-protein interaction (Andrade and Bork, 1995; Takano and Gusella, 2002), leading to the proposal that HTT is a scaffold to mediate protein interactions and integrate multiplex cellular responses (Cattaneo et al., 2005; MacDonald, 2003). Indeed, HTT is involved in a diverse set of cellular processes including transcription, trafficking and autophagy (Liu and Zeitlin, 2017; Saudou and Humbert, 2016). However, despite extensive studies, it is still unclear how HTT itself is regulated *in vivo*.

As a powerful genetic model, *Drosophila* has been invaluable in discovering and dissecting conserved signaling pathways and modeling human diseases including HD (Xu et al., 2015). Previously, we have characterized *Drosophila* HTT homolog (*dhtt*) (Zhang et al., 2009), which at amino acid level has relatively low homology with human HTT (Li et al., 1999). But resembling HTT, *dhtt* also encodes a large protein (3,511 amino acids) consist primarily of predicted HEAT repeat (Li et al., 1999; Takano and Gusella, 2002; Zhang et al., 2009). Importantly, beyond their similarity in large protein size and secondary structure, *dhtt* is also involved in similar cellular processes as its mammalian counterparts. For example, *dhtt* can functionally replace HTT in rescuing the axonal trafficking and misaligned mitotic spindle in mouse cortical neurons (Godin et al., 2010; Zala et al., 2013) and plays a similar conserved role in selective autophagy (Ochaba et al., 2014; Rui et al., 2015), supporting *Drosophila* as a relevant *in vivo* organism for functional dissection of HTT.

Considering the structural and functional conservation of HTT from flies to humans, we hypothesize that the core regulators of HTT, especially those with close physical interactions with HTT, should be conserved between these two evolutionary distant species. Indeed, among the large number of HTT associated proteins (HAPs) identified from various biochemical and proteomic studies, most have fly homologues (Goehler et al., 2004; Harjes and Wanker, 2003; Kaltenbach et al., 2007; Li et al., 2007; Saudou and Humbert, 2016; Shirasaki et al., 2012; Tourette et al., 2014). However, by now few of them have been validated *in vivo*, and the central regulators of HTT remain largely obscure.

Accordingly, we carried out proteomic studies to directly isolate dHtt-associated proteins (dHaps) from fly tissues, from which we identified a novel 40kDa protein as a specific and potentially strongest binding partner of dHtt. This 40kDa protein, encoded by a previous uncharacterized gene *cg8134*, shares notable sequence similarity with HAP40, a known HTT binding partner in mammals. HAP40 was first isolated from rat brain homogenate as an interactor of endogenous HTT (Peters and Ross, 2001), and was shown later in a proteomic study in mouse brains as the “*most significantly correlated*” with HTT in their protein levels (Shirasaki et al., 2012). Consistently, HAP40 was found to bind HTT tightly at 1:1 molar ratio (Pal et al., 2006). More recently, HAP40 was found to be important in regulating HTT’s conformation (Guo et al., 2018). Importantly, a ∼10 fold increase of HAP40 levels were observed in primary fibroblasts and striatal tissues from postmortem brains as compared to healthy controls (Pal et al., 2006). Despite these strong biochemical and structural evidence, there is no reported study on HAP40’s physiological roles in any animal settings, and its functional relationship with HTT and its potential involvement in HD pathogenesis remain unclear.

We found that *in vitro*, the 40kDa protein encoded by *cg8134* was capable of physically interacting with human HTT. Conversely, when expressed in flies, human HAP40 could fully rescue the phenotypes of *cg8134-*null mutants, supporting *cg8134* as the fly orthologue of HAP40. Accordingly, we renamed *cg8134* as *dhap40* (*Drosophila* Hap40). At whole animal levels, flies lacking *dhap40* showed similar loss of function phenotypes as *dhtt* null (*dhtt-ko*) mutants, suggesting that *dhtt* and *dhap40* play similar or overlapping physiological roles. Strikingly, there was an almost complete loss of endogenous dHap40 protein in *dhtt* null flies. Similarly, the levels of endogenous dHtt were significantly reduced in *dhap40* knockout flies. Importantly, a similar mutual reliance between HTT and HAP40 is conserved in human cells. Lastly, when tested with fly models of HD, HAP40 selectively modulated neurodegeneration induced by full-length mutant HTT but not mutated HTT exon 1, potentially through its specific effects both on the overall protein levels and on the toxicity of mutant HTT *per se*. Collectively, these findings demonstrate that HAP40 is an essential and highly conserved regulator of HTT *in vivo* and a potential therapeutic target for HD.

## Results

### Affinity purification of interacting partners for endogenous dHtt from *Drosophila*

We first examined the expression patterns of endogenous dHtt protein by Western blot analysis, which revealed a single protein product of about 400kDa that was present in wildtype but absent in *dhtt-ko* flies. Resembling the mammalian expression pattern, dHtt protein was widely expressed at low levels at all developmental stages and tissues examined, with relatively high levels in early stage embryos and in adult brains (Fig. 1A and data not shown).

**Figure 1.**
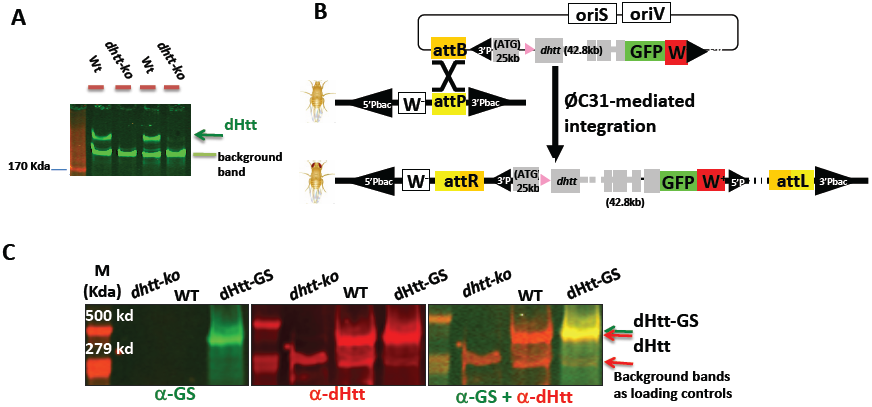
dHtt expression and genomic tagging of dHtt. (A) Western blot analysis of endogenous dHtt expression in early stage *Drosophila* embryos, revealed with anti-dHtt antibody. A single ∼400kDa dHtt band is present in wildtype embryos and absent in age-matched *dhtt-ko* mutants. The lower background band served as loading control, as noted. (B) Schematics for genome-tagging *dhtt* gene at its C-terminus with eGFP, which includes an ∼80kb genome region covering *dhtt* gene, including about 25kb upstream of the first coding exon, 42.8 kb genome region covering all the coding exons and about 12kb 3’-untranslated region (UTR). eGFP was fused in-frame to the C-terminus of encoded dHtt protein before the stop codon and 3’ UTR. The genome fusion constructs were cloned into pacman vector through the recombineering method, and subsequently integrated into the preselected attP site in the fly genome through the phiC31-integrase mediated transgene approach (see Methods). (C) Western analysis for the expression of dHtt-GS-TAP fusion protein from pacman-*dhtt-GS-TAP* transgenic flies (dHtt-GS), or controls of wildtype (WT) and *dhtt-ko* mutants, as indicated. The GS-TAP tag contains two Protein G modules, a TEV protease cleavage site, and a streptavidin binding peptide (SBP). The whole-protein extracts from the indicated genotypes were probed simultaneously with anti-SBP (green band in left panel) to detect the GS-TAP tag and anti-dHtt (red bands in the middle panel) antibodies. The dHtt-GS-TAP fusion protein was only detected in the *dhtt*-GS transgenic flies and absent in controls of WT (middle lane) and *dhtt-ko* (left lane) flies. The overlaying image (right panel) for both anti-SBP and anti-dHtt staining showed that the dHtt-GS-TAP was expressed as full-length fusion protein (orange band) with slightly larger size than endogenous dHtt. A background band of ∼300kDa size served as loading control.

Given its overall low expression levels, to achieve effective pulldown of its binding partners under physiological conditions, we created two genome-tagging lines containing in-frame fusion of eGFP or GS-TAP tags at the C-terminus of the encoded dHtt protein. Specific anti-GFP antibodies, including the nanobody (Cristea et al., 2005), are available for efficient pull down of GFP-tagged protein, while GS-TAP is a tag optimized for tandem-affinity-purification (TAP) of target proteins from *Drosophila* tissues or cultured cells (Burckstummer et al., 2006; Kyriakakis et al., 2008). The GFP and GS-TAP tags have no overlap sequences, thus allowing affinity-purification (AP) of dHtt and its associated proteins (dHaps) via two complementary approaches. Using pacman, an optimized BAC transgene approach (Venken et al., 2006), we established the genome-tagging lines for *dhtt-eGFP* and *dhtt-GS-TAP* (Fig. 1B). The ∼83Kb BAC constructs carried the full coding and regulatory regions of *dhtt* gene, thereby enabling the expression of the tagged *dhtt* transgene under the control of its native regulation elements, and the levels and expression patterns of the tagged dHtt protein mirroring that of endogenous dHtt. Subsequent characterization of the pacman transgenic flies confirmed the expression of the tagged dHtt-eGFP and dHtt-GS-TAP proteins at the expected sizes and levels comparable to that of endogenous dHtt (Fig. 1C and data not shown). Further, despite their relatively low abundance in whole animal homogenates, dHtt-GS-TAP fusion was significantly enriched to relatively high purity through the TAP sequential purification (Fig. 2A and 2B). Similarly, a single-step pulldown by anti-GFP nanobody highly enriched dHtt-GFP from homogenates (Fig. 2C and data not shown). The *dhtt* genome tagging lines thus allow convenient isolation of dHaps under physiological setting, avoiding artifacts often associated with protein mis-expression and overexpression.

**Fig 2.**
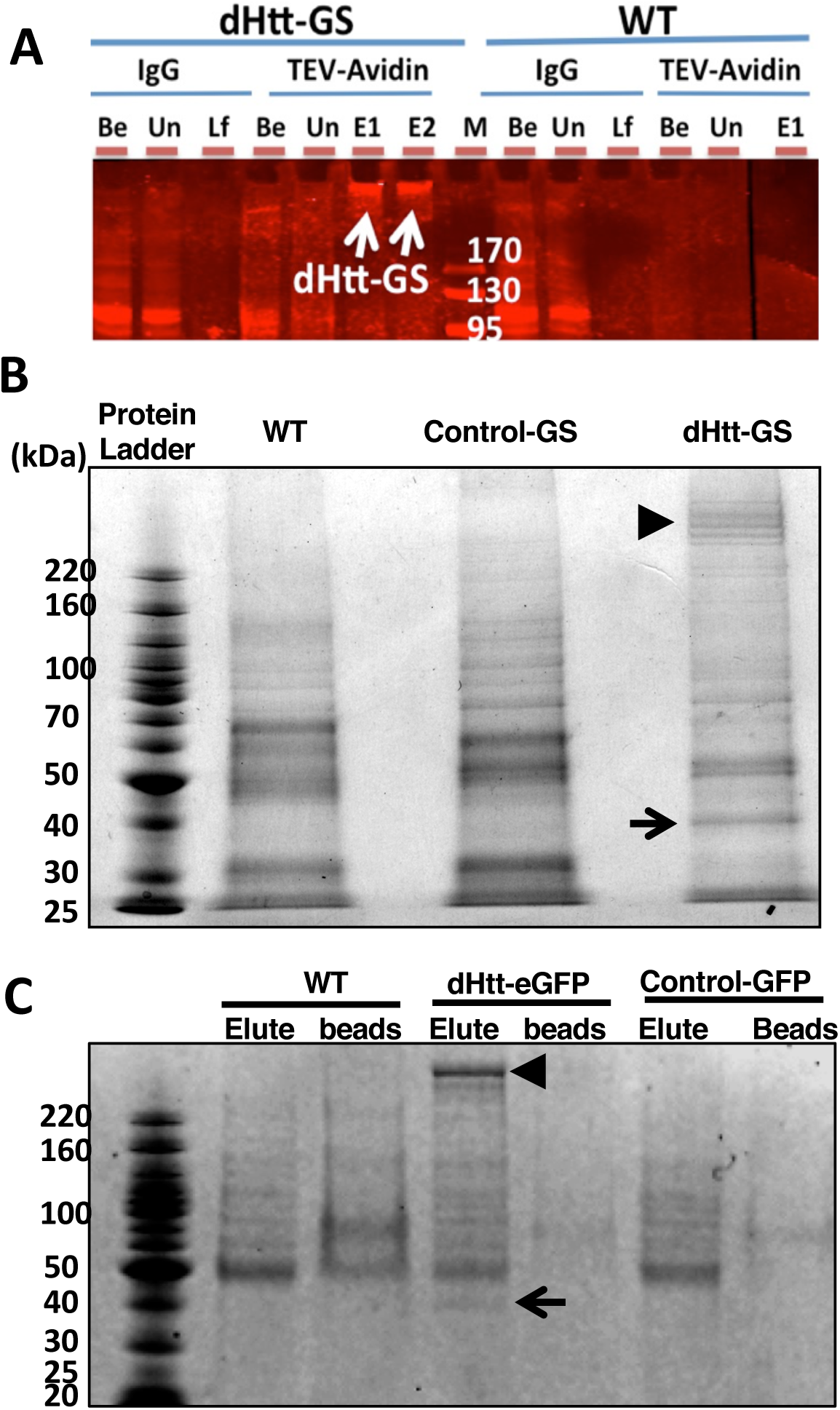
Affinity-purification of endogenous dHtt and dHtt-associated proteins from *Drosophila*. (A) Western blot analysis of sequential purification steps for dHtt-GS-TAP fusion and its associated proteins from flies. Whole embryos extracts from wildtype control (WT) or flies transgenic for pacman-dhtt-GS-TAP (dHtt-GS) were processed in parallel through the following purification steps: (1) IgG-pulldown against Protein G modules in the GS-TAP tag; (2) TEV protease cleavage of the TEV recognition site in the GS-TAP tag to release dHtt and associated proteins from IgG beads; (3) Second pull-down with Streptavidin beads against the SBP within the GS-TAP tag; (4) final elution of dHtt and dHtt-associated proteins from Streptavidin beads with biotin. When possible, equal volumes of protein extracts or agarose slurry from the following purification steps were analyzed, as annotated in the figure panel (from the left to right): Be, whole embryos extract *before* incubation with IgG beads; Un, *unbound* protein samples after depletion of dHtt-GS with IgG-conjugated agarose beads from whole embryos extracts; Lf, *leftovers* on IgG beads after TEV cleavage; Be, elute from IgG beads after TEV cleavage, *before* incubating with Streptavidin-conjugated agarose beads; E1 and E2: final *elute* 1 and 2 fractions released from Streptavidin beads with biotin solutions. Samples from each of the above purification steps were processed for SDS-PAGE analyses and probed with rabbit α-dHtt antibody. Note that the quantity of endogenous dHtt protein in the crude protein extracts from the pacman-dhtt-GS-TAP transgenic flies was barely detectable (lane 1, labeled as “Be” in left panel), and became significantly enriched in the final elutes (arrows in E1, E2). The same band was absent from WT control (right panel). M: protein ladder *marker*, with their sizes labeled. (B and C) SAS-PAGE analysis of affinity-purified dHtt and dHtt-associated proteins isolated from transgenic flies carrying (B) pacman-*dhtt*-GS-TAP (dHtt-GS), or (C) pacman-*dhtt*-eGFP, visualized with Coomassie blue staining. The following controls were processed in parallel: wildtype (WT) non-transgenic flies; controls for pacman flies carrying in-frame fusion with an unrelated protein of (B) GS-TAP (Control-GS) or (C) eGFP (Control-GFP) tags, as indicated. Note the co-purification of a prominent ∼40kDa protein (arrows) with the ∼400 kDa dHtt (arrowheads) specifically from (B) dHtt-GS or (C) dHtt-GFP flies only, but not from either of the controls. In (B), several large bands were present around the ∼400 kDa range (arrowhead) specifically in dHtt-GS sample, which likely were partially degraded dHtt protein generated during multi-step TAP purification procedures. In (C), after final eluting from GFP-nanobody agarose beads with 10mM glycine (PH 2.5), both the elutes (Elute) and the post-elution agarose beads (beads) were processed for SDS-PAGE analysis.

### CG8134 is a strong and specific binding partner of endogenous dHtt

Using *dhtt* genome-tagging flies, we optimized and then carried out multiple rounds of large-scale affinity purifications for dHtt associated proteins from whole embryo homogenates. To eliminate false positives common with purification procedures or with promiscuous binding to the GS-TAP or GFP tags, we included three sets of controls in parallel experiments: (1) wildtype flies carrying no transgenes, (2) an independent pacman transgenic line expressing an unrelated gene with a C-terminal GS-TAP tag, and (3) another pacman transgenic line expressing the same unrelated gene with a C-terminal GFP tag.

In all the pull-down assays, a prominent ∼40kDa band was co-purified with both dHtt-eGFP and dHtt-GS-TAP, but not in either of the controls (arrows in Fig. 2B and 2C). The relatively high abundance of this ∼40kDa protein in independent dHtt-pull down experiments suggests that it is a specific and likely the strongest binding partner for endogenous dHtt. Further, considering that dHtt is almost ∼10 times larger in size than this ∼40kDa protein, the relative signal densities of the co-purified dHtt and the ∼40kDa bands indicates that these two proteins might complex at about 1:1 molar ratio (Fig. 2C and data not shown). Mass spectrometry of the 40kDa band revealed it is the predicted protein product of a previously uncharacterized gene *cg8134* (Fig. 3A).

**Fig 3.**
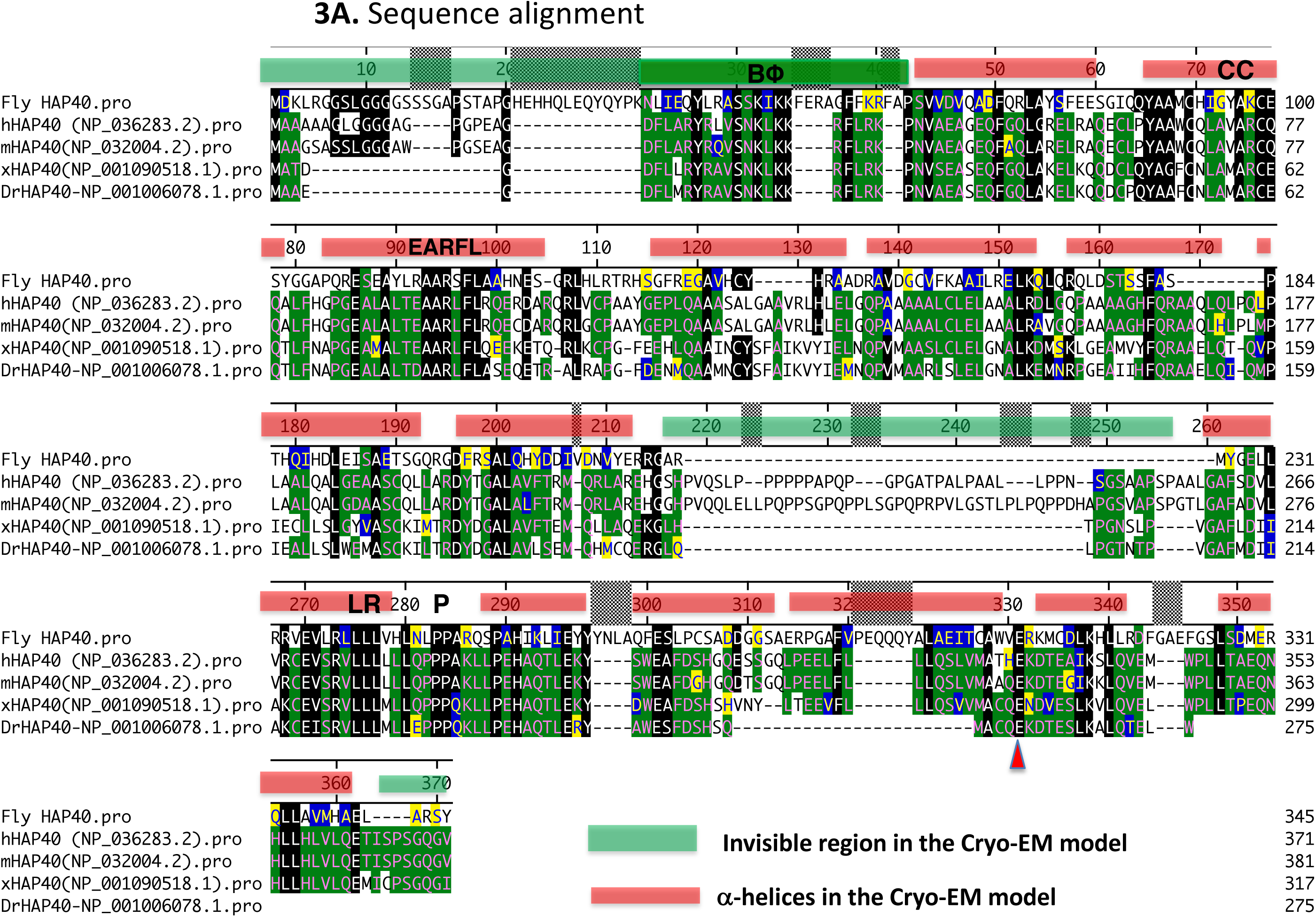
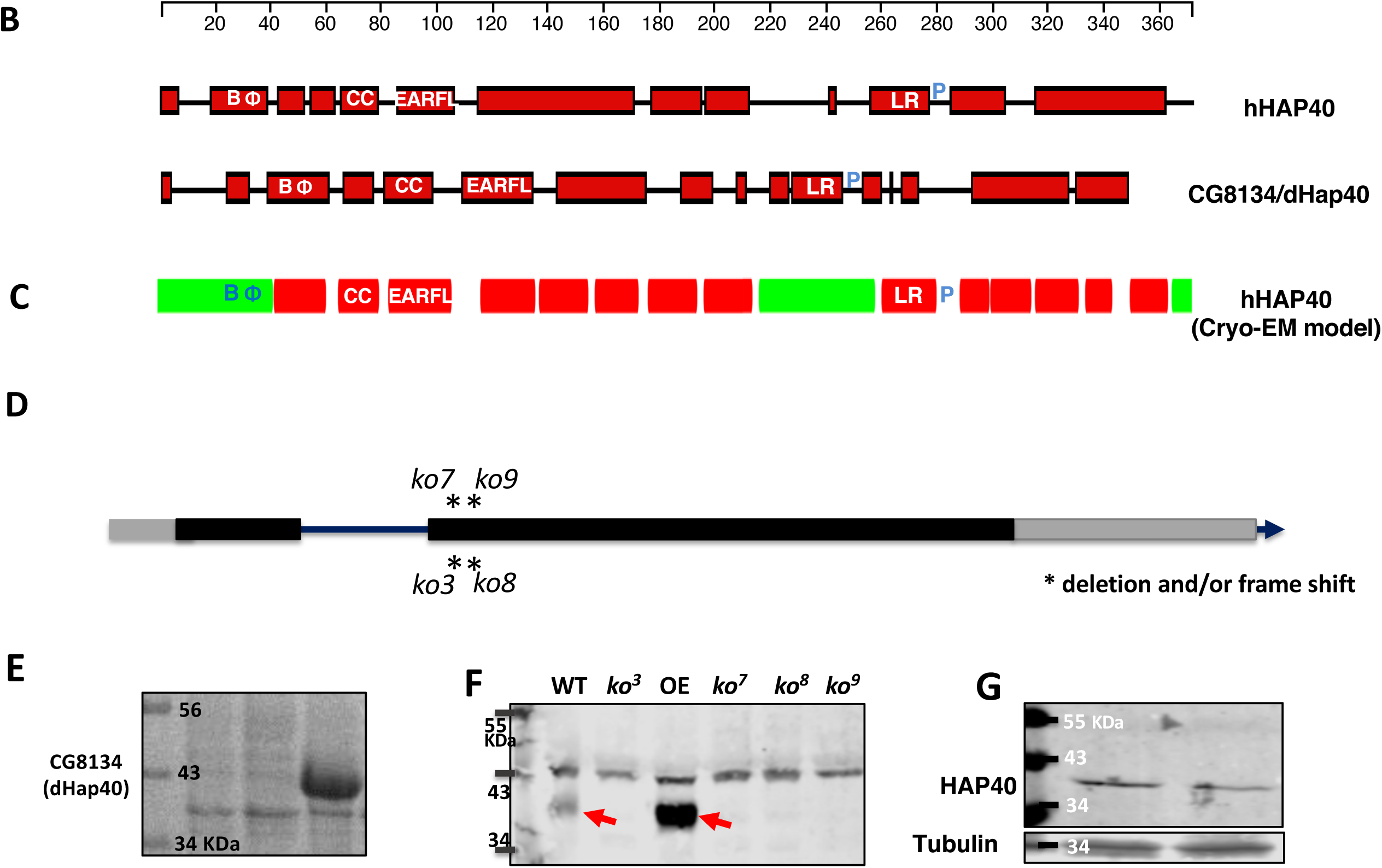
CG8134 encodes *Drosophila* Hap40 homologue. (A) Sequence alignment between CG8134/dHap40 and the following vertebrate HAP40 homologs: human (hHAP40/F8a1), mouse (mHAP40/F8a), xenopus (xHAP40) and zebra fish *Danio rerio* (DrHAP40). Genebank ID of these proteins were indicated. The scale bar above the alignment refers to amino acid position from human HAP40 (NP_036283). Amino acid similarities are annotated in color in the following order in the alignment: (1) black box highlights amino acids that are identical to that in CG8134/dHap40; (2) green boxes are those identical to those in human HAP40; (3) Yellow boxes are those with similar chemical property as those in human HAP40; (4) blue box highlights amino acids that are with similar chemical property as in CG8134/dHap40. The red bars above the alignment cover the sequences of the predicted 14 α-helices and the green bars those invisible in the Cryo-EM model. (B) Schematics of the predicted secondary structures in human HAP40 (top) and CG8134/dHap40 (bottom) proteins using the Garnier-Robson structure prediction module (Protean program by DNASTAR), drawn in scale (top) of human HAP40. Red boxes represent the predicted α-helices. BΦ: the N-terminal α-helices enriched with basic and hydrophobic amino acids, which is part of the N-terminal invisible region in the Cryo-EM model (see Fig. 3C below); CC: the α-helices containing two conserved cysteines; EARFL: α-helices with a conserved stretch of E,A,R,F and L amino acids; LR: α-helices with leucine-repeat; P: proline linker after the LR α-helices. (C) Schematics of the secondary structures in human HAP40 protein predicted from the Cryo-EM study. Red rectangles correspond to the predicted 14 α-helices and the green rectangles are regions invisible in the Cryo-EM model. (D) Schematics of genomic structure of *cg8134/ dhap40* gene and the mutant alleles created in the study. *dhap40* is a X-linked gene composed of one intron (solid line) and two coding exons (grey boxes: untranslated regions; black boxes: coding regions), with arrowhead indicting the orientation of the encoded protein from 5’ to 3’ ends. Stars annotate the locations of the molecular lesions for each of the characterized *dhap40* alleles. (E) SDS-PAGE and Coomassie blue staining of protein lysates from *E. coli.* A protein product of 40kDa size was produced only in cells transformed with a plasmid containing full-length *cg8134* cDNA (lane 3) but not in controls transformed with an empty protein expression vector alone (lanes 1 and 2). (F) Western Blot analysis of CG8134/dHap40 expression in adult flies from wildtype (WT), four *dhap40* mutant (*ko3, ko7, ko8* and *ko9*) alleles, and a dHap40 overexpression line (OE) carrying a UAS-*cg8134* transgene directed by a strong Actin-Gal4 driver. Note that a 40kDa band (left red arrow) corresponding to endogenous CG81344/dHap40 was detected in WT and present at significantly higher levels in the OE line (right red arrow), but was completely absent in the four *dhap40* mutant lines. (G) Western Blot analysis of ectopically human HAP40, detected as a 40kDa protein, from homogenates of two fly lines carrying a UAS-F8A1/HAP40 transgene direct by *daughterless*-Gal4 driver.

### CG8134 encodes the fly HAP40 ortholog

BLAST search of the protein encoded by *cg8134* revealed notable similarity with HAP40 homologs from multiple vertebrate species (Fig. 3A). The recent Cryo/EM study on the structure of the HTT/HAP40 complex noted that homology-based search failed to identify a HAP40 homolog in *Drosophila* (Guo et al., 2018). However, the homology between CG8134 and HAP40 exist throughout most of the proteins, and are particularly prominent in two ∼50 amino acid (a.a.) long stretches at their N- or C-terminal regions, where there are about ∼40% identical and ∼50% similar amino acids between CG8134 and human HAP40 proteins (Fig. 3A and Supplemental Fig. S1). Importantly, the Cryo/EM study revealed that the N- and C-regions of HAP40 mediate its direct physical interactions with HTT (Guo et al., 2018), raising an attractive possibility that the similar conserved regions in CG8134 also mediate its binding with dHtt in *Drosophila*.

The homology is not just limited to amino acid sequence, but also their predicted secondary structures. In particular, when analyzed using the same modeling parameters, both CG8134 and human HAP40 are predicted to be composed mainly of α-helices that are distributed in similar patterns throughout the two proteins (Fig. 3B). Importantly, these predicted α-helices also align remarkably well with the 14 α-helices identified in the Cryo-EM study (Guo et al., 2018).

Sequence alignment across species also revealed several intriguing structural features suggestive of potential functional importance (Fig. 3A-C). For example, the central region of human HAP40, predicted as unstructured from the sequence-based prediction (Fig. 3B), is invisible in the model from the Cryo-EM study (the middle green bar in Fig. 3C) and absent in both CG8134 and the HAP40 homologs in zebrafish and frogs (Fig. 3A), consistent with the findings that it is dispensable for HTT binding (Guo et al., 2018). In contrast, although the very N-terminus of HAP40 is also invisible in the Cryo-EM study, it contains a ∼20 amino acid-long stretch which we named as “BΦ” motif for its unusually high content of basic and hydrophobic residues, a feature that is highly conserved (Fig. 3A-C). In particular, in human HAP40, the BΦ motif contains eight strong basic (K,R) and eight hydrophobic (A,I,L,F,W,V) amino acids (a.a. 21-41, DFLARYRLVSNKLKKRFLRKP), while the corresponding stretch in CG8134 has seven strongly basic and nine hydrophobic amino acids (a.a. 38-61, NLIEQYLRASSKIKKFERAGFFKR). Such a unique and conserved amino acid composition implies a potential functional importance for this N-terminus motif. Similarly, the second predicted α-helices (labeled as “CC” in Fig. 3A-C) is characterized by two highly conserved cysteine residues (a.a. 70 and 76 in human HAP40. Fig. 3A). The predicted ninth α-helices is marked by its long leucine repeat (“LR” in Fig. 3A-C, a.a. 260-280 in human HAP40), containing four continuous leucine residues in dHap40 and six leucine in human and mouse HAP40, followed by two or three continuous proline (“P” in Fig. 3A-C) before the next α-helices. Lastly, E331 at the C-terminus of human HAP40 exist across all the species (red arrowheads in Fig. 3A), supporting it functional importance in mediating the direct interaction with the bridge domain of HTT as revealed in the Cryo-EM study (Guo et al., 2018).

When expressed in *E. coli*., *cg8134* indeed produced a protein product at the predicted ∼40kDa size (Fig. 3E). Moreover, in whole animal homogenates, two independent antibodies raised against CG8134 both detected a ∼40kDa band that was present in wildtype flies but completely absent in all the established *cg8134*-null mutant lines (Fig. 3D and 3F, see below). Further, in flies with ectopic overexpression of *cg8134* from a UAS-*cg8134* transgene, which served as a positive control for CG8134 expression, the levels of the same ∼40kDa band became significantly higher (Fig. 3F). Lastly, when human HAP40 protein was expressed in fly tissues from transgenic flies, it also ran at similar size as fly *cg8134* (Fig. 3G). Taken together, these results demonstrated that the ∼40kDa protein co-purified with dHtt protein is the protein product of *cg8134* gene. The overall sequence and structural similarities between CG8134 and vertebrate HAP40 suggest that *cg8134* encodes the fly ortholog of HAP40, a conclusion supported in subsequent functional studies (see below). We therefore renamed the *cg8134* gene as *dhap40* (*Drosophila* HAP40 (HTT-associated protein 40kDa)).

### *dhap40* and *dhtt* share similar phenotypes in *Drosophila* and have similar conserved roles in autophagy

To investigate the physiology functions of *dhap40* in *Drosophila*, we generated UAS-based transgenic flies for *cg8134* full-length cDNA that allow its targeted overexpression in different fly tissues (Fig. 3F)(Brand and Perrimon, 1993), and also created multiple independent *cg8134* mutant alleles using the CRISPR/Cas9 approach (Bassett et al., 2013; Cong et al., 2013; Gratz et al., 2013; Port and Bullock, 2016) (Fig. 3D and Supplemental Fig. S2). The established *dhap40* mutant alleles harbor various molecular lesions, including deletions and frame shifts, in the coding region of *dhap40* gene (Fig. 3D and Supplemental Fig. S2). Consistently, for each of the alleles, the endogenous dHap40 protein was absent in homogenates of homozygous mutant flies (Fig. 3F), suggesting that they are all protein-null. Notably, all these established *dhap40* null alleles are homozygous viable and developed normally into adulthood. Further, both male and female mutant flies are fertile with no apparent morphological defects (data not shown). These observations are reminiscent that of *dhtt*-null flies, which are also homozygous viable and fertile with no apparent developmental defects (Zhang et al., 2009). However, the *dhtt*-null adults develop ageing-related phenotypes, including shortened lifespan, reduced mobility and mild autophagy defects (Ochaba et al., 2014; Rui et al., 2015; Zhang et al., 2009). Remarkably, *dhap40-ko* mutants manifested very similar phenotypes as *dhtt* null. All the *dhap40-ko* alleles examined died early, with an average life span of 40 days, compared to 33 days for *dhtt*-null flies and 59 days for WT flies (Fig. 4A). As the flies become older, the *dhap40-ko* flies also showed accelerated reduction of their mobility, as measured by climbing assay, mirroring that of *dhtt*-null flies (Fig. 4B and data not shown). Importantly, all observed phenotypes could be rescued by ectopically expressed *dhap40* from the UAS-*dhap40* (*cg8134*) transgenes driven by a ubiquitously *daughterless*-Gal4 (da-Gal4), thus confirming that the observed defects of *dhap40-ko* animals are due to the loss of endogenous dHap40 (Fig. 4C and data not shown). The similarity in loss of function phenotypes between *dhtt* and *dhap40* flies suggest that the two genes share similar physiological roles and likely function together in the same or overlapping cellular processes. However, it is noticeable that overall the phenotypes of *dhap40* mutants were relatively weaker than that of *dhtt-ko* flies, living on average about 7 days longer (Fig. 4A) and their mobility declining at a slower pace (Fig. 4B), potentially indicating a modulatory role by HAP40 within the HTT/HAP40 complex in facilitating the cellular functions executed by HTT.

**Fig 4.**
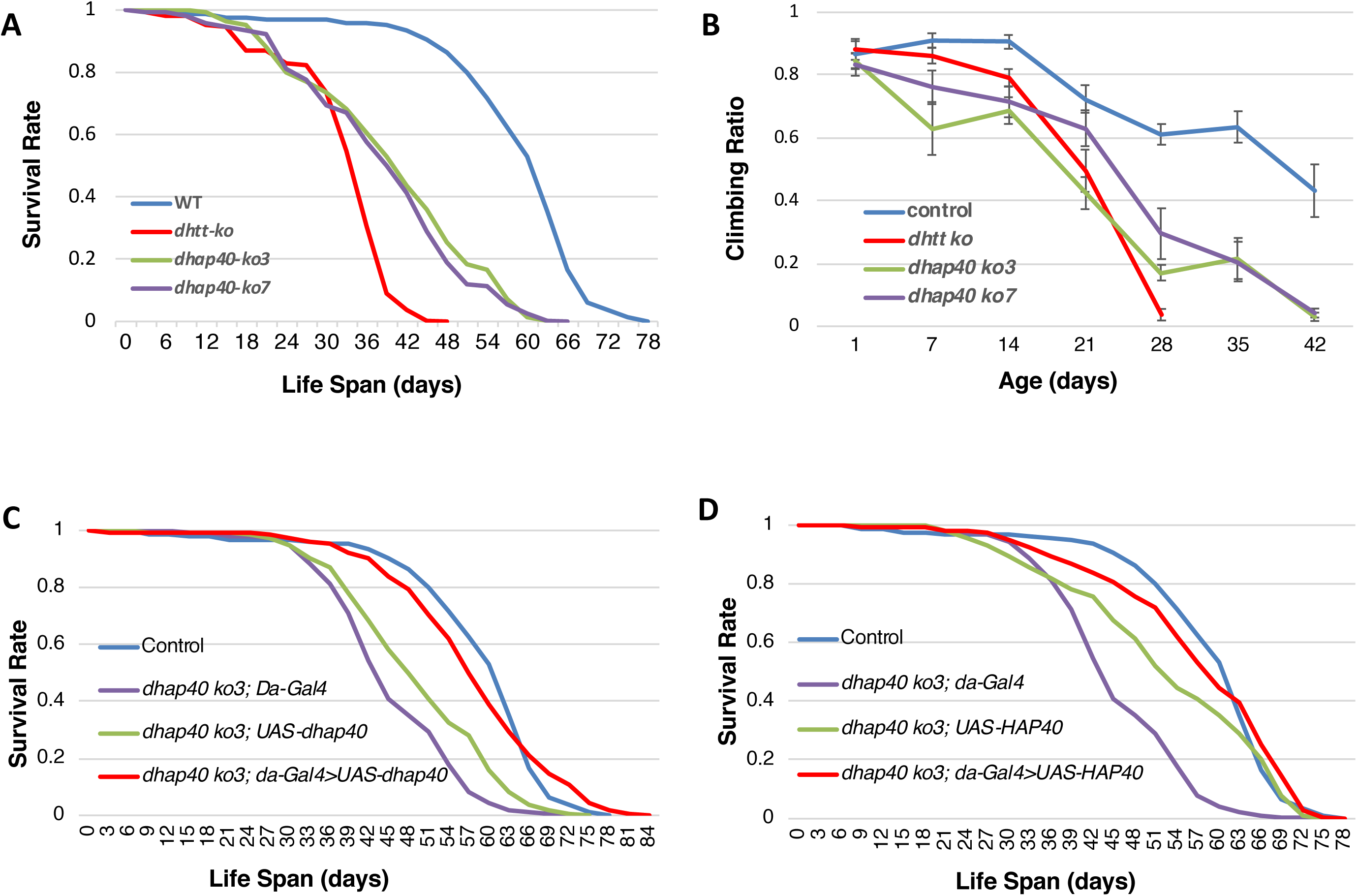
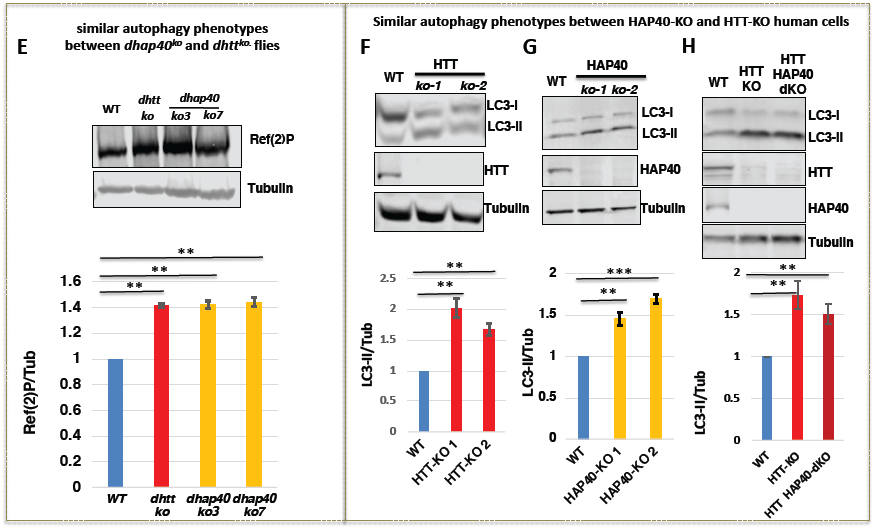
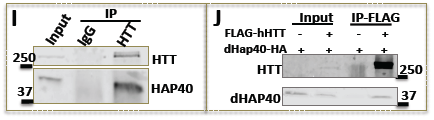
*dhap40* mutants show similar loss-of-functions phenotypes as *dhtt* null flies. (A) Survival curves and (B) climbing assays for two *dhap40* null alleles *ko3* and *ko7* (green and purple lines, respectively), together with wildtype control (blue line) and null *dhtt-ko* mutants (red line), as indicated. Note that both *dhap40* mutant alleles manifested similar, albeit significant weaker, phenotypes than *dhtt-ko* flies. (A) *dhap40*^ko^ had an average life span of 40 days (n=230 total), differing significantly (p<0.001, log-rank test) from both wildtype (59 days, n=253) and null *dhtt-ko* mutants (33 days, n=209). (B) Climbing assays for 28-day-old adult flies of the indicated genotypes, with *dhap40* flies (n=180) showing significant differences (p<0.01 in both cases, t-test) from both wildtype (n= 200) and *dhtt-ko* mutants (n=180). (C and D) Rescue experiment for the longevity deficit of *dhap40*^*ko3*^ flies, restored by ectopically expressed (C) fly dHap40 from the UAS-dhap40 (*cg8134*) or (D) human HAP40 from UAS-HAP40 (F8a1) transgenes, all directed by a ubiquitous *da*-Gal4 driver (red lines. Genotype: “*w*^*1118*^, *dhap40*^*ko3*^; UAS-*cg8134*/da-Gal4” (C, n=300) or “*w*^*1118*^, *dhap40*^*ko3*^; UAS-F8a1/da-Gal4” (D, n=307)). The survival curves of the rescued flies in (C) and (D) were statistically indistinguishable from wildtype (blue lines, p>0.5, log-rank test n=253. Genotype: “*w*^*1118*^”), but significantly different from the two controls: (1) *dhap40*^*ko3*^ flies carrying da-Gal4 driver alone (purple lines. p<0.001, log-rank test. Genotypes: “*w*^*1118*^, *dhap40*^*ko3*^; da-Gal4/+”. n=308); and (2) *dhap40*^*ko3*^ flies carrying (C) UAS-*dhap40* (Genotype: “*w*^*1118*^, *dhap40*^*ko3*^; UAS-*cg8134*/+”. n=355) or (D) UAS-HAP40 (Genotype: “*w*^*1118*^, *dhap40*^*ko3*^; UAS-F8a1/+”. n=337) transgene alone (green lines, p<0.001, log-rank test), as indicated. The same set of data from wildtype (blue lines) and “*dhap40*^*ko3*^; da-Gal4/+” (purple lines) were used in both (C) and (D) as shared controls. (E-H) HAP40 and HTT share similar autophagy defects in (E) flies and (F-H) human cells. (E) Western blot assays for Ref(2)p in whole protein extracts from adult flies of two *dhap40* null alleles (*ko3* and *ko7*) together with *dhtt-ko* null or wildtype (WT) controls, as indicated. (F-H) Western blot assays for LC3-II in whole cell lysates from wildtype (WT) controls and (F) two independent HTT-KO HeLa cell lines, (G) two HAP40-KO HEK293 cell lines, or (H) HTT-KO and HTT/HAP40-dKO HeLa cell lines, as indicated. Bar charts below show quantification results of normalized (E) Ref(2)P or (F-H) LC3-II levels from three independent repeat experiments for each assays. ** p< 0.05. *** p< 0.01 (student’s t-test). α-Tubulin served as loading control in all experiments. (I and J) co-IP experiments between (I) endogenous HTT and HAP40 or between (J) transfected HTT and dHap40 in HEK293 cells, as indicated. Note that the pulldown efficiency by HTT was significantly higher against (I) endogenous HAP40 than (J) dHap40.

At the cellular level, one of the established roles of HTT is in autophagy. Consistently, resembling that observed in *dhtt-ko* flies (Ochaba et al., 2014; Rui et al., 2015), loss of *dhap40* led to increased accumulation of Ref(2)p, the fly homologue of a known autophagy substrate p62/SQSTM (Fig. 4E and data not shown)(Nezis et al., 2008). To test if HAP40 has a similar conserved role in mammalian cells, we next created multiple independent HTT knockout (HTT-KO) and HAP40 knockout (HAP40-KO) lines from HEK293 cells, which were validated in Western blot assays for the absence of the endogenous HTT and HAP40 proteins, respectively (Fig. 4F-H). Both HTT-KO and HAP40-KO cell lines were viable and behaved similarly in subsequent assays. Resembling the HTT-KO cells (Fig. 4F and data not shown), HAP40-KO cells showed a similar increase of lipidated LC3 (LC3-II), a marker for the autophagosome. Similar autophagy defects were also observed in HTT-KO and HAP40-KO lines derived from HeLa cells, supporting that the role of HAP40 in autophagy regulation is not cell-type specific (Fig. 4G and data not shown).

Although HTT and HAP40 are likely to act together in a single protein complex in autophagy regulation, it is still possible that HTT and HAP40 effect autophagy through different or parallel pathways. To exclude this possibility, we next created a stable HeLa cell line with double knockout (dKO) for both HTT and HAP40 genes. Western blot assay confirmed the simultaneous loss of both HTT and HAP40 proteins in this dKO line (Fig. 4H). Again, the HTT/HAP40 double-KO cells showed comparable levels of autophagy defects as in either single HTT- or HAP40-knockout alone (Fig. 4H and data not shown), supporting that HTT and HAP40 function together as a single complex and play similar physiological role in autophagy regulation.

### HAP40 is functionally conserved from flies to humans

The above results prompted us to examine the potential functional conservation of HAP40 between the fly and humans. We first tested whether human HAP40 could rescue the loss of function phenotypes of *dhap40-ko* flies. Indeed, when directed by the same da-Gal4 driver, human HAP40 significantly suppressed all the observed mutant phenotypes of *dhap40-ko* flies, including the shortened lifespan and reduced mobility (Fig. 4D and data not shown). Such a functional rescue would predict that *Drosophila* dHtt still retains the ability to physically interact with human HAP40 protein, and dHap40 with HTT. Indeed, in HEK293 cells, HTT could co-immunoprecipitate (co-IP) ectopically expressed dHap40 protein from transfected cells (Fig. 4J), although the pulldown efficiency was significantly lower than the robust pulldown of endogenous HAP40 by HTT in parallel co-IP experiments (Fig. 4I). This conserved physical interaction is consistent with the observation that the N- and C-terminal regions that are important for HTT-binding are highly conserved between dHap40 and HAP40 (Fig. 3A and Supplemental Fig. S1), which might be the underlying basis for the functional conservation of HAP40 between these two evolutionarily very distant species. Together, these results demonstrate that *Drosophila* CG8134/dHap40 is the functional ortholog of HAP40.

### High levels of HAP40 is not toxic in *Drosophila*

An early study reported an up to ∼10-fold increase of the levels of HAP40 protein in samples from HD patients and mouse HD models, potentially implicating a toxic effect from elevated HAP40 expression (Pal et al., 2006). We therefore examined the effects of HAP40 overexpression in various fly tissues. Using the established UAS-transgenic lines, driven by different Gal4 lines, we achieved dHap40 overexpression at levels ∼10 folds or more higher than that of the endogenous dHap40 (Fig. 3F). However, in all the Gal4 lines tested, including the ubiquitous (i.e., da-Gal4, Act-Gal4 and Tubulin-Ga4), neuronal-specific (i.e., Elav-Gal4 and Syt-Gal4) and eye-specific GMR-Gal4, dHap40 overexpression did not cause any apparent effect such as animal lethality and eye degeneration (data not shown). Similarly, no apparent detrimental effects were observed in flies overexpressing human HAP40 from the established transgenic lines (Fig. 3G and data not shown). Together, they suggest that high levels of HAP40 might not be toxic by itself, at least in the tested *Drosophila* tissues.

### HTT and HAP40 are mutually dependent for their protein stability

Given the highly conserved physical and functional interaction between HTT and HAP40, we next tested how they might regulate each other. Strikingly, in *Drosophila*, in two independent null *dhtt* alleles, loss of dHtt led to an almost complete depletion of endogenous dHap40 protein (Fig. 5A). Importantly, such regulation is also conserved in mammalian cells, as in independent HTT-KO cell lines derived either from HEK293 or HeLa cells, the levels of endogenous HAP40 were dramatically reduced to about ∼10-20% of that in control cells (Fig. 5B and data not shown). Importantly, restoring HTT expression in the HTT-KO cells, by transfecting plasmids encoding full-length HTT, led to a full rescue of the levels of endogenous HAP40 protein. Moreover, increased expression of ectopic HTT similarly resulted in higher levels of endogenous HAP40 (Fig. 5C and data not shown).

**Fig 5.**
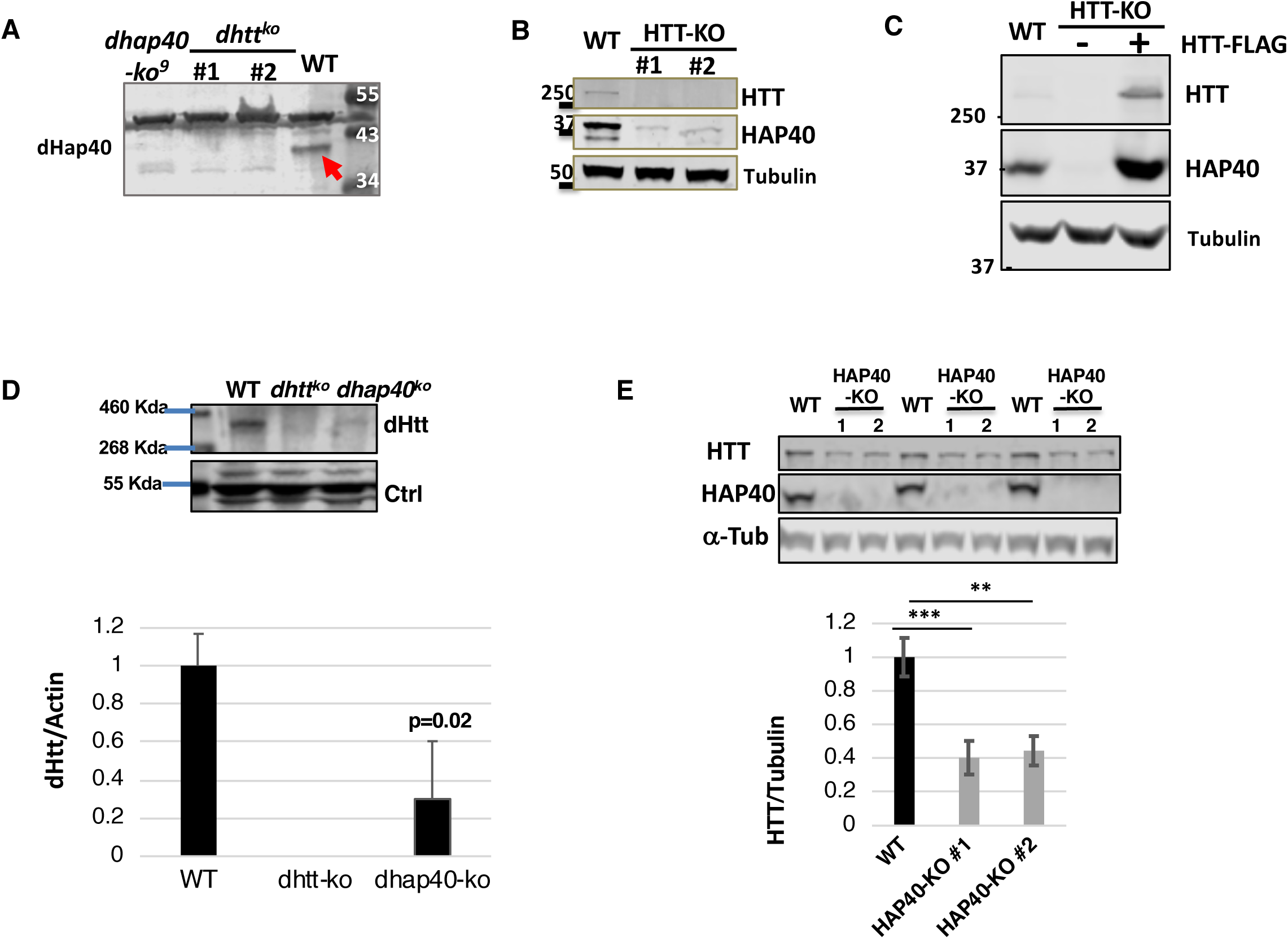
Conserved mutual dependence between HAP40 and HTT proteins on each other’s stability. Western blot assays of endogenous HTT and HAP40 proteins in flies and human cells. (A) The 40kDa endogenous dHap40 (arrow) protein in wildtype control (WT) was absent in two independent *dhtt* null alleles, similar as a *dhap40* null allele. Genotypes as indicated. A background band of 50kDa size present in all samples served as loading control. (B) Endogenous HAP40 was significantly depleted in two independent HTT-knockout (HTT-KO) HEK293 cell lines, as compared to wildtype HEK293 control (WT). (C) The levels of HAP40 was severely depleted in HTT-KO cells (lane 2) and was restored by overexpressed FLAG-HTT to a level even higher (lane 3) than in WT control (lane 1). (D) Reduced levels of endogenous dHtt protein in null *dhap40* ^*ko3*^ adult flies as compared to WT and *dhtt*^*ko*^ controls, with an average of 70% reduction in *dhap40*^ko^ flies as compared to WT control shown in the bar chart below of the quantification from three repeat experiments (p=0.02, student’s t-test). Actin (not shown) and ∼50kDa background bands from anti-dHap40 antibody served as loading controls. (E) Reduced levels of endogenous HTT protein, shown in three independent Western blot assays, in two independent HAP40-KO cell lines and their quantification in the bar chart below, showing an average of ∼60% reduction of normalized HTT levels as compared to WT control. ** p< 0.05. *** p< 0.01 (student’s t-test). α-Tubulin serves as loading and normalization controls in all the panels unless otherwise indicated.

Given the above findings, we next examined whether HTT levels are similarly affected by the loss of HAP40. In *Drosophila*, in all *dhap40-ko* alleles we tested, there were a significant, ∼70% reduction of the levels of endogenous dHtt, in comparison with that in wildtype controls (Fig. 5D). Similarly, in both human HEK293T and HeLa cells, loss of HAP40 led to a notable ∼60% reduction of the levels of endogenous HTT protein (Fig. 5E). Together, these results revealed that HTT and HAP40 proteins are mutually dependent on each other for their own *in vivo* stability, a regulation that is highly conserved during evolution from flies to humans.

### HTT and HAP40 have different protein dynamics

We further explored the mechanism underlying the mutual dependence between HTT and HAP40 proteins. In HTT-KO cells, application of proteasome inhibitor MG132 partially restored the levels of endogenous HAP40. In contrast, autophagy and lysosome inhibitors such as chloroquine (CQ), Bafilomycin A1 (BafA1) or ammonium had no discernable effect on the levels of endogenous HAP40 (Fig. 6A, quantification in Fig. 6B and data not shown), suggesting that in the absence of HTT, HAP40 is quickly cleared by the proteasome system. However, when similarly applied to HAP40-KO cells, neither of the tested proteasome or lysosome inhibitors could significantly rescue the diminished levels of HTT protein within the duration of the drug treatment (Fig. 6C, quantification in Fig. 6D and data not shown). Together, these results suggest very different protein dynamics between HTT and HAP40, with the smaller HAP40 protein being a preferential target of the proteasome in the absence of HTT, while the much larger HTT being relatively more stable and refractory to degradation in the absence of HAP40.

**Fig 6.**
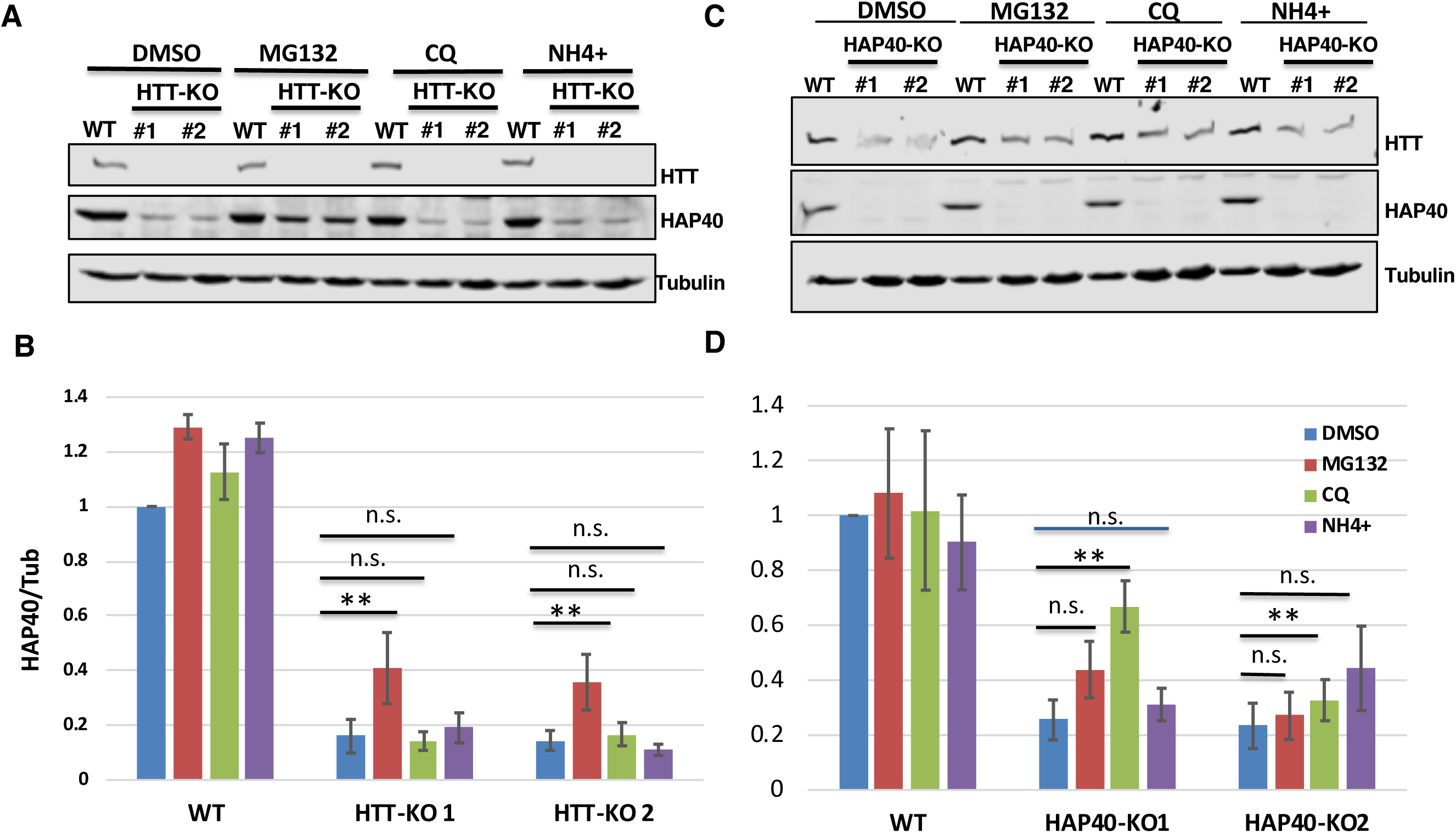
Proteasome mediates HAP40 degradation. Western blot assays and quantifications for endogenous HAP40 and HTT proteins in HTT-KO, HAP40-KO or wildtype HEK293 (WT) cells under different treatments, as indicated. (B and D) Normalized levels of (B) HAP40 or (D) HTT proteins from three repeat experiments, corresponding to (A) and (C), respectively. Treatment with proteasome inhibitor MG132 for 5 hours partially but significantly restored the levels of depleted HAP40 protein in HTT-KO cells (A and B), but showed no clear effect on the levels of endogenous HTT protein in HAP40-KO cells (C and D). Autophagy/lysosome inhibitors ammonium (NH4+) and CQ, also treated for 5 hours, behaved similarly as DMSO mock treatment in both HTT-KO and HAP40-KO cells. ** p< 0.05 (student’s t-test). n.s., no significance. α-Tubulin served as loading and normalization controls in all the experiments.

### PolyQ expansion in HTT has minimal effect on its binding affinity for HAP40

As polyQ length correlates with mutant HTT toxicity, we next examined whether polyQ expansion in HTT could affect its affinity for HAP40. We first performed co-IP assays between HAP40 and FLAG-tagged full-length HTT proteins carrying a polyQ track with 23, 72 or 145 glutamines. In transfected HEK293T cells, wildtype and mutant HTT proteins showed comparable pull-down efficiency against endogenous HAP40, quantified based on the ratios of co-immunoprecipitated HAP40 and HTT proteins in repeat experiments (Fig. 7A and quantification in Fig. 7B). Interestingly, albeit not statistically significant, the quantification result implied a trend of slightly higher affinity between HAP40 and mutant HTT-145Q, whereas the binding affinity of HAP40 with HTT-23Q and HTT-73Q were virtually the same (Fig. 7B). To further test the above observation, we took advantage of the findings that the stability of endogenous HAP40 relies almost exclusively on the presence of HTT (Fig. 5), and compared the efficiency of wildtype and mutant HTT in restoring the levels of endogenous HAP40 in HTT-KO HEK293 cells. Quantification data from this rescue experiment showed that in HTT-KO cells, both mutant HTT-73Q and HTT-145Q restored the levels of endogenous HAP40 protein as effectively as wildtype HTT-23Q (Fig. 7C and quantification in Fig. 7D). Nevertheless, time course analyses, by comparing the relative levels of HAP40 protein versus that of the ectopically expressed HTT at different time points after HTT transfection in HTT-KO cells, again suggested a trend of slightly better rescue efficiency by HTT-145Q than HTT-73Q and HTT-23Q in restoring the expression of endogenous HAP40 (Figure 7D), although the difference was too weak to reach a statistical significance in this HAP40 rescue experiment. Lastly, as ectopic expression of HTT in wildtype cells can induce higher levels of endogenous HAP40 (Fig. 5C), we further compared wildtype or mutant HTT for their overexpressing effect on the levels of endogenous HAP40 protein in normal HEK293T cells. The results were similar as that in HTT-KO cells, with HTT-145Q being slightly more effective than HTT-23Q and HTT-73Q in inducing higher levels of endogenous HAP40 (data not shown). Taken together, these data suggest that polyQ expansion in HTT does not compromise, and might even mildly enhance, its affinity for HAP40.

**Fig 7.**
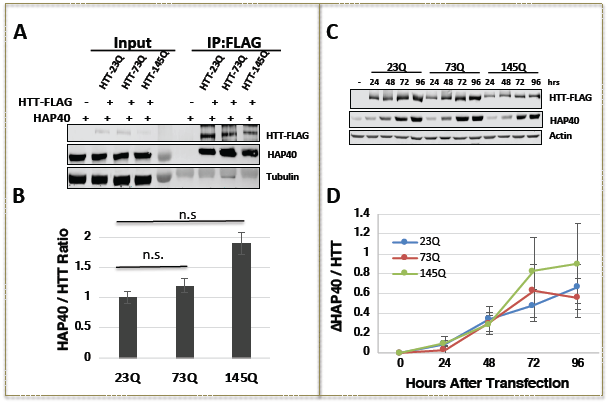
PolyQ expansion in HTT does not significantly affect its affinity for HAP40. (A) Western blot assay and (B) quantification of co-IP experiments between HAP40 and FLAG-tagged full-length HTT carrying 23, 72 or 145 (23Q, 72Q and 145Q) glutamine tracts, as indicated. (B) The pull-down efficiency of HAP40 by different HTT proteins were quantified as the relative ratio of co-immunoprecipitated HAP40 and HTT proteins, averaged from three independent experiments. All three HTT proteins showed comparable pulldown efficiency for endogenous HAP40. n.s., no significance. α-Tubulin serves as loading control. (C and D) Time course analyses of endogenous HAP40 and ectopically expressed HTT-23Q, 72Q or 145Q in HTT-KO HEK293 cells, hours after transfection with FLAG-tagged HTT expressing plasmids, as indicated. (D) Quantification of the time-dependent changes of HAP40 levels normalized against HTT at each time point, averaged from three repeat experiments.

### HAP40 is a specific toxicity modulator of full-length mutant HTT

Considering the close physical and functional relationship between HAP40 and HTT, one important question is whether and how HAP40 might modulate the toxicity of mutant HTT. Multiple *Drosophila* models of HD have been created that can faithfully recapitulate the polyQ length-dependent toxicity of mutant HTT proteins (Xu et al., 2015), thus allowing us to address this question in this model organism. We first tested flies with neuronal expression of human HTT exon1 carrying 93 glutamines (HTTex1-93Q), a well-established HD model that manifests robust neuronal degeneration in targeted neurons (Steffan et al., 2001). Interestingly, loss of endogenous *dhap40* did nor apparently modify the neuronal degeneration of HTTex1-93Q flies, showing a similar loss of internal photoreceptor cells in *dhap40-ko* flies as in wildtype background (Supplemental Fig. 3).

Given that HAP40 binds full-length HTT (fl-HTT), but not its N-terminal region alone (Guo et al., 2018; Pal et al., 2006), we next tested another well-characterized HD model expressing mutant human fl-HTT with 128Q (fl-HTT-128Q) or wildtype control of fl-HTT with 16Q (fl-HTT-16Q) (Romero et al., 2008). In addition, to ensure the reproducibility and minimize background variation, we also created a new set of UAS-based transgenic lines for human fl-HTT carrying 23, 73 or 145 glutamines, with all the lines being inserted to the same attP40 integration site in the fly genome through the attB/Phi31 targeted integration method (Bischof et al., 2007; Groth et al., 2004). As expected, when tested using different Gal4 drivers, including the eye-specific GMR-Gal4 and pan-neuronal *elav*-Gal4 or *syt*-Gal4 drivers, wildtype fl-HTT (16Q or 23Q) did not induce any discernable detrimental effect, fl-HTT-73Q caused an intermediate whereas fl-HTT-128Q and fl-HTT-145Q much stronger toxicity, including early animal lethality and prominent eye degeneration, the latter being manifested as adult-onset gradual depigmentation and loss of internal photoreceptor cells (Fig. 8A-D and data not shown). When tested using the full-length HTT-based HD models, loss of *dhap40* modestly ameliorated the neurodegenerative phenotypes. For example, loss of *dhap40* suppressed the depigmentation phenotype of flies with eye-specific expression of fl-HTT-145Q (compare Fig. 8B with 8A), as well as the degeneration of internal neuronal photoreceptor cells of the flies with pan-neuronal expression of fl-HTT-128Q (Fig. 8C). Consistently, the adult lethality linked to pan-neuronal expression of mutant HTT was also mildly but clearly delayed in the absence of *dhap40* (Fig. 8D).

**Fig. 8.**
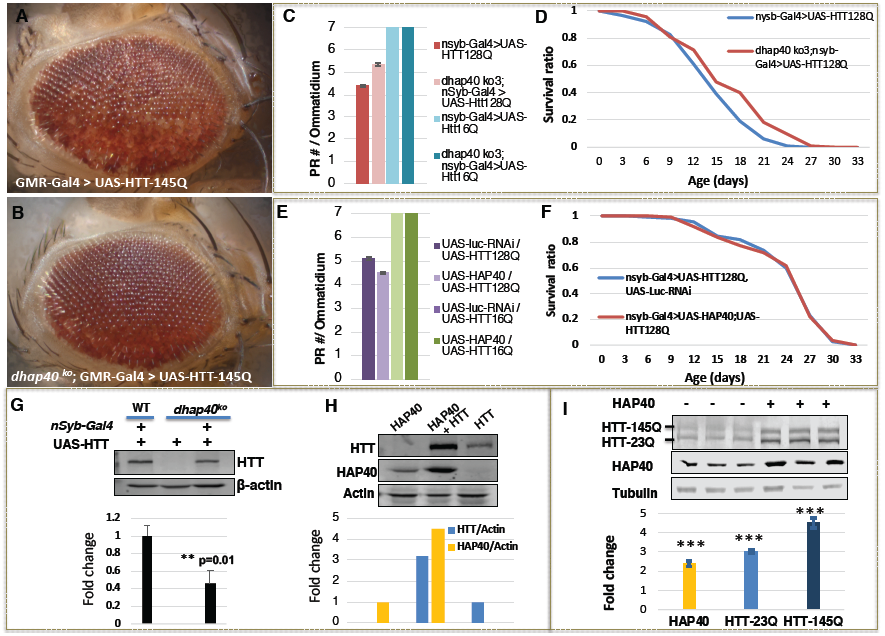
HAP40 modulates neurodegeneration of *Drosophila* HD model through multiple mechanisms. (A and B) Bright-field image of 30-day-old adult fly eyes expressing fl-HTT-145Q driven by eye-specific GMR-Gal4 in (A) wildtype and (B) *dhap40*^ko3^ background. The loss of eye pigmentation induced by fl-HTT-145Q (A), indicating underlying eye degeneration, was suppressed by the absence of *dhap40* (B). (C-E) Quantification of photoreceptor cell degeneration and viability phenotypes of flies with neuronal-specific expression of fl-HTT-128Q either (C and D) in *dhap40* null background or (E and F) with co-expression of human HAP40, all directed by pan-neuronal *nsyb*-Gal4 and with respective controls, as indicated. (C and E) Bar chart presentation of the average number of intact photoreceptor cells (PRC) per ommatidium in 7-day-old flies of the following designated genotypes. (C)*”nsyb-*Gal4/UAS*-*fl-HTT-128Q*”*: 4.4 PRC/ommatidium (n=360 ommatidia from 20 flies); “*dhap40* ^ko3^; *nsyb-*Gal4*/*UAS*-*fl-HTT-128Q”: 5.3 PRC/ommatidium (n=360 ommatidia from 20 flies); The difference between the two was significance (p<0.001, Mann-Whitney rank-sum test). Control flies expressing Fl-HTT-16Q in both normal and *dhap40*-ko backgrounds showed normal seven photoreceptor cells (blue bars). (E) “UAS*-*HAP40; *nsyb-*Gal4 *>*UAS*-*fl-HTT-128Q” flies: 5.1 PRC/ommatidium (n=331 ommatidia from 20 flies); control “UAS*-*Luciferase-dsRNA; *nsyb-*Gal4 *>*UAS*-*fl-HTT-128Q” flies: 4.5 PRC/ommatidium (n=320 ommatidia from 20 flies). The difference between the two is significance (p<0.001 by Mann-Whitney rank-sum test). Control flies co-expressing HTT-16Q with HAP40 or with luciferase (green bars) all had seven intact photoreceptor cells. (D and F) Survival curves of the adult flies expressing fl-HTT-128Q. (D) Average life span was 17 days for “*dhap40* ^*ko3*^; *nsyb-*Gal4*/*UAS*-*fl-HTT-128Q” flies (n=179) and 15 days for control *“nsyb-*Gal4*/*UAS*-*fl-HTT-128Q*”* flies (n=229). The difference between the two is significance (p<0.001. Log-rank test). (F) Average life span was 24.3 days for flies co-expressing HTT-Q128 with HAP40 (genotype: “*nsyb*-Gal4>UAS-HAP40/+; UAS-HTT-128Q/+”. n=141), and 24.5 days for control flies co-expressing HTT-Q128 with luciferase-dsRNA (“*nsyb*-Gal4> UAS-luciferase-RNAi/+; UAS-HTT-128Q/+”, n=174), with no significant difference between the two genotypes (p=0.9 by Log-rank test). (G-I) Western blot assays for the ectopically expressed human HTT protein in (G and H) flies and (I) human cells, and quantifications from three independent repeat experiments show in bar charts below. (G) The levels of human HTT expressed from the same UAS-HTT transgene was ∼60% lower in *dhap40-ko* (lane 3) than in normal (WT, lane 1) background, and was absent in control flies lacking *nSyb*-Gal4 driver (lane 2). (H) The levels of HTT were about four times higher and HAP40 about three times higher in flies co-expressing HTT and HAP40 (lane 2) than flies expressing HTT (lane 3) or HAP40 (lane 1) alone, all driven by *nSyb*-Gal4. (I) Western blot assays for HTT and HAP40 levels in HEK293T cells with simultaneous co-expression of FLAG-tagged HTT-23Q and HTT-145Q, ran as three repeat experiments probed with anti-FLAG and anti-HAP40 antibodies, as indicated. Co-transfection with HAP40 led to an average of about two-fold increase of HAP40 and three- to five-fold increase of HTT-23Q and HTT-145Q (compare lane 4-6 with lanes 1-3). *** p< 0.001 (student’s t-test), which were calculated as the fold changes in HAP40 and HTT levels after co-transfection with HAP40 as compared to before HAP40 transfection.

The weak but significant protective effect conferred by the loss of endogenous *dhap40* would predict that higher levels of HAP40 should exacerbate the neurodegeneration of HD flies. Surprisingly, co-expression of human HAP40 only marginally enhanced the eye degeneration of the fl-HTT-128Q-expressing flies (Fig. 8E). Moreover, the animal lethality associated with mutant fl-HTT was literally unchanged in the presence of the co-expressed HAP40 (Fig. 8F). As expected, co-expression of HAP40 with wildtype HTT (fl-HTT-23Q or fl-HTT-16Q) did not elicit any apparent phenotypes in all the tissues examined (Fig. 8C, 8E and data not shown).

In different animal models of neurodegenerative diseases, the severity of phenotypes mostly correlates with the expression levels of mutated disease proteins such as mutant HTT (Romero et al., 2008; Warrick et al., 1998; Yamamoto et al., 2000). Therefore, the observed modifications by endogenous *dhap40* or the ectopically expressed human HAP40 could be due to their effect on overall HTT protein levels, especially considering the strong and conserved mutual dependence between HTT and HAP40 in maintaining each other’s stability (Fig. 5). Accordingly, we examined whether the levels of human HTT, expressed ectopically from the transgenic flies, could be affected by the loss of endogenous *dhap40* or by the presence of co-expressed human HAP40. Indeed, when the same UAS-HTT transgene line was driven either by pan-neuronal *syt*-Gal4 or other drivers, the levels of HTT in *dhap40-ko* mutants was less than half of that in wildtype background (Fig. 8G and data not shown). Conversely, co-expression of human HAP40 resulted in over three-fold increase of HTT levels than when HTT was expressed alone and strikingly, a similar increase of co-expressed HAP40 than when HAP40 was expressed alone (Fig. 8H and data not shown). These results implied that when expressed ectopically in transgenic flies, human HTT can be stabilized not only by human HAP40, but also by endogenous fly dHap40, an observation in line with the findings that fly dHap40 can bind, albeit at reduced efficiency, with human HTT (Fig. 4J). Consistently, when tested in human cells, a similar synergistic effect was also observed between co-expressed HTT and HAP40. For example, in HEK293T cells that simultaneously co-expressed wildtype (23Q) and mutant (145Q) fl-HTT proteins, co-transfection of HAP40 led to significantly increased levels of both fl-HTT-23Q and fl-HTT-145 HTT proteins (compare lanes 4-6 with lanes 1-3 in Fig. 8I and the quantification chart below).

Collectively, the above results reveal a rather selective and complex role of HAP40 on HTT-induced neurodegeneration. First, HAP40’s effect on mutant HTT toxicity is more specific to full-length HTT, not truncated HTT such as HTT exon 1, a finding that is consistent with the observation that HAP40 primarily interacts with full-length HTT (Guo et al., 2018; Pal et al., 2006). Second, HAP40 can influence neurodegeneration through its profound effect on HTT’s stability and total protein levels, as loss of endogenous *dhap40* halved the levels of ectopically expressed HTT protein (Fig. 8G) and correspondingly mildly suppressed the neurodegeneration phenotypes (Fig. 8A-D). This finding also implies that HAP40-free mutant HTT largely retains it toxicity, if not more toxic. However, the effect on HTT levels alone can not be the sole mechanism on how HAP40 modulates mutant HTT-induced neurodegeneration, as it cannot explain the observation that co-expression of HAP40 resulted in over three-folds increase of total HTT levels (Fig. 8H) but only a very marginal or negligible (Fig. 8E-F) effect on the phenotypes of the tested HD flies. Rather, they raise an intriguing possibility that HAP40 plays double-edged roles in HD, with HAP40-binding leads to stabilized HTT and its significantly elevated levels, but also converts mutant HTT to a less toxic form, one that harbors lower potency than HAP40-free HTT in inducing neurodegeneration when present at equal molarity, thereby neutralizing the exacerbating effect normally associated with higher levels of mutant HTT protein.

## Discussion

Despite the evidence implicating the importance of HAP40 on HTT regulation and HD pathogenesis, there has been little systematic examination of HAP40 in physiological and pathological settings. From a forward proteomic study in *Drosophila*, we isolated a novel 40kDa CG8134 protein as one of the strongest binding partners of the HTT homolog in *Drosophila* and further demonstrated it is the fly ortholog of HAP40, with conserved physical and functional interactions with HTT. The co-existence of HAP40 and HTT in evolutionarily distant species from flies to humans not only supports the functional importance of HAP40 in HTT biology, but also establishes *Drosophila* as a relevant genetic model to evaluate the physiological and pathological roles of HAP40. Further studies demonstrated that HAP40 is a central and positive regulator of HTT’s endogenous functions, and there exists a strong mutual-dependence between endogenous HTT and HAP40 on each other’s stability, a regulation that is conserved from flies to humans. Lastly, glutamine expansion in HTT does not significantly affect its affinity for HAP40, while HAP40 can potentially be an important modulator of HD pathogenesis through its multiplex effect on HTT’s stability and overall protein levels, its physiological functions, and the potency of mutant HTT toxicity *per se*.

### HAP40 is a conserved binding partner of HTT

In the recent Cryo-EM study on HTT structure, it was noted that a homology-based search failed to identify a HAP40 homolog in the *Drosophila* genome (Guo et al., 2018). This is probably due to the overall low degree of sequence similarity between human HAP40 and fly dHap40, mirroring the significant sequence divergence between human and fly HTT from two very distant species (Li et al., 1999). However, our results clearly demonstrated that the previously uncharacterized *cg8134* gene encodes the fly orthologue of HAP40, as evident by its tight association with endogenous dHtt protein from an *in vivo* pull-down assay (Fig. 2), by their overall structural similarities, especially their highly conserved N- and C-terminal regions known to be important for HTT binding (Fig. 3), by the retained ability of dHap40 to physically interact with human HTT (Fig. 4J) and stabilize the ectopically expressed HTT in flies (Fig. 8G), and lastly, by the ability of human HAP40 to rescue the loss of function phenotypes of *dhap40*-null flies (Fig. 4D).

The importance of this conserved partnership between HTT and HAP40 is further reinforced by the findings of a highly conserved mutual dependence on each other’s stability between the two proteins. In both flies and human cells, loss of dHtt or HTT leads to an almost complete depletion of endogenous dHap40 or HAP40, respectively (Fig. 5). Conversely, higher levels of HTT resulted in elevated levels of HAP40 (Fig. 5C); and both in flies and human cells, their co-expression led to a synergistic, multi-fold increase of both proteins than when each was expressed alone (Fig. 8H and 8I). These findings together support that complex formation with HAP40 is an important stabilizing mechanism for HTT. Collectively, our findings not only establish *Drosophila* as a relevant animal model for studying HAP40, but also uncover an ancient and important regulatory mechanism governing HTT that constrains the co-evolution of this HTT/HAP40 partnership through millions of years of evolution pressure in flies and humans.

The similarity between dHap40 and vertebrate HAP40 proteins exist at both sequence and structural levels (Fig. 3A-C), which not only form the basis of their functional conservation, but also implicate physiological significance of the conserved regions in HAP40 proteins. For example, the N- and C-terminal regions as well as residue E331 of human HAP40 are all highly conserved from *Drosophila* to vertebrates, supporting their roles in mediating direct physical interactions with HTT (Guo et al., 2018). Conversely, consistent with its dispensable role in HTT binding, the central region of human HAP40 is not conserved in fly dHap40 and coincidentally is also invisible in the Cryo-EM imaging (Guo et al., 2018). Among the other conserved regions in HAP40 (Fig. 3A-3C), the 20 a.a.-long BΦ motif at the N-terminus is particularly interesting, as it is invisible in the Cryo-EM model, indicating its structural flexibility; however its highly conserved and rather unique amino acid composition potentially suggest an important functional role. Similarly, the invariable proline repeats in-between the predicted α-helices 9 and 10 (Fig. 3A-3C. prolines 282 to 284 in human HAP40) potentially introduce an important structural kink that facilitates the proper folding of HAP40 proteins. As an extrapolation, the size of HAP40 proteins is also relatively conserved, at around ∼40kDa. Since within the HTT/HAP40 complex, HAP40 is sandwiched inside HTT (Guo et al., 2018), this raises an intriguing possibility that HAP40 in its folded conformation needs to stay within specific size range in order to fit in and prop up HTT into a unique shape or conformation, so as to properly execute its cellular functions. Future structural-functional study should help clarify the physiological significance of these observed conservations.

Our results also indicate that in the absence of HTT, most of HAP40 is degraded quickly through the proteasomal machinery (Figs. 5 and 6). Conversely, in both *dhap40*-null flies and HAP40-knockout human cells, lower but significant levels of HTT protein persist (Fig. 5), suggesting very different turnover dynamics of these two proteins.

Finally, many studies on HTT and HD primarily focus on N-terminal HTT fragments, especially its exon 1, which lack the ability to physically interact with HAP40 (Guo et al., 2018; Pal et al., 2006). Given that endogenous HTT mainly exists in a complex with HAP40 while HAP40-free HTT is functionally or activity-wise quite different from HTT/HAP40 complex (Figure 4), the HAP40-binding capacity of HTT constructs should be considered in future functional studies.

### HAP40 is a positive regulator of HTT’s endogenous functions

HTT has diverse cellular functions from transcription to autophagy (Liu and Zeitlin, 2017; Saudou and Humbert, 2016). Given that HAP40 is an essential HTT partner, it raises questions on whether HAP40 positively or negatively affects HTT’s cellular activities, and whether the two proteins always function together in the same processes or each mediates certain specific cellular activities. Identification of the HAP40 homolog in *Drosophila* affords us the opportunity to address these questions. Our studies in flies and human cells support that HAP40 is a conserved positive regulator of HTT. In *Drosophila, dhap40*-null flies manifest similar phenotypes as *dhtt-ko* flies, including compromised autophagy as well as reduced mobility and shortened lifespan (Fig. 4A-E). Consistently, HAP40-KO human cells exhibited similar autophagy defects as HTT-KO cells (Fig. 4F-H). Further, human cells with HTT and HAP40 double-KO show similar autophagy defects as HTT or HAP40 single knockout alone (Fig. 4H). Nevertheless, in *Drosophila*, the overall phenotypes of *dhap40* null flies are weaker than that of *dhtt-ko* mutants, living on average 7 days longer (Fig. 4A) and mobility declines slower than *dhtt-ko* flies (Fig. 4B). Although loss of HAP40 markedly reduced levels of endogenous HTT, significant amounts of HTT remained in both flies and human cells (Fig. 5), which would suggest that the HAP40-free HTT is still partially functional, or alternatively they might mediate a distinct set of cellular functions independent of the HTT-HAP40 complex. Collectively, our findings support that HTT and HAP40 largely function together in the same set of cellular processes, in which HAP40 primarily acts as a positive regulator of HTT. Further, the two proteins are not functionally equal, as the primary role of HAP40 is likely to stabilize and regulate HTT, a hypothesis consistent with the observations that loss of HTT leads to an almost complete depletion of endogenous HAP40 in flies and human cells, while HTT alone is more stable and likely still partially functional.

### HAP40 modulates HD pathogenesis through its multiplex effect on full-length HTT

An early study reported an almost ten-fold increase of the levels of HAP40 in samples from HD patients and mouse models (Pal et al., 2006). However, when expressed in different fly tissues, either alone or together with wildtype human HTT, higher levels of fly dHap40 or human HAP40 did not cause apparent detrimental effect, suggesting that elevated levels of HAP40 by itself might not be toxic. Considering that *in vivo*, the majority, if not all, of HAP40 protein should be in complex with HTT as most HTT-free HAP40 is cleared away by the proteasome (Figs. 5 and 6), it raises another question on how HAP40 becomes stabilized and accumulates to abnormally high levels under pathological conditions, especially giving the findings that in HD brains, the levels of mutant HTT protein are similar as or even lower than that of wildtype HTT from the normal allele (Aronin et al., 1995; Bhide et al., 1996). One plausible explanation might be that HAP40 in mutant HTT complex is more resistant to degradation, as suggested by the early observation that mutant HTT can interfere with the proteasomal machinery (Seo et al., 2004; Wang et al., 2008), which we show is involved in HAP40 clearance (Fig 6).

As an essential regulator of HTT, some other key questions are how HAP40 affects the toxicity of mutant HTT, and whether the HAP40-free mutant HTT is more or less toxic. We found that polyQ expansion in HTT did not effect, but only marginally increased, its binding with HAP40 (Fig. 7). This is consistent with the findings that mutant HTT largely retains its key physiological functions (Hodgson et al., 1996; White et al., 1997) and accordingly its ability to associate with its major binding partners including HAP40. However, our results support that HAP40 behaves as a specific and important regulator of HD pathogenesis through its multiplex effect on full-length HTT. Most telling, *in vivo*, when expressed from the same transgene line, the levels of mutant full-length HTT in *dhap40*-null background was significantly lower than in wildtype (Fig. 8G), but induced rather comparable degrees of neurodegeneration (Fig. 8A-D). Considering the close correlation between the levels of mutant proteins and the severity of the neurodegeneration in animal models of neurodegenerative diseases including HD in flies (Romero et al., 2008; Warrick et al., 1998; Yamamoto et al., 2000), these data not only indicate that the HAP40-free mutant HTT is still toxic, but when present at equal molar amount, HAP40-free mutant HTT might be even more potent in its toxicity than mutant HTT in complex with HAP40. Consistent with this hypothesis, co-expression of human HAP40 resulted in multi-fold increase of HTT protein levels, but only a rather marginal enhancement of eye degeneration phenotypes (Fig. 8E) and virtually no effect on animal survival in the tested HD flies (Fig. 8F). Together, these findings lead to our conclusion that HAP40-free mutant HTT is less stable but more potent in its toxicity, while HAP40-binding significantly stabilizes mutant HTT but also reduces its potency of toxicity.

These observations suggest that HAP40 might profoundly affect HD pathogenesis through multiple avenues, by governing the physiological functions of HTT, by controlling HTT protein’s stability and thereby its overall levels, and by modulating the potency of mutant HTT toxicity *per se*. First, in animal models of neurodegeneration, toxicity of mutant HTT often correlate with the levels of the expressed protein (Romero et al., 2008; Warrick et al., 1998; Yamamoto et al., 2000). We demonstrate that HAP40 is essential for HTT’s stability and overall protein levels (Fig. 5), with HTT levels greatly elevated with total levels of HAP40 in targeted fly tissues and mammalian cells, and loss of endogenous dHap40 reduced the amount of ectopically expressed human HTT (Fig. 8G-I). The latter observation (Fig 8G) is consistent with the finding that dHap40 retains a conserved despite weaker binding ability with human HTT (Fig 4J). Second, normal functions of HTT are known to be neuronal protective, potentially arising from its roles from autophagy to trafficking and neurotrophic effect by BDNF and other neurotrophic factors (Liu and Zeitlin, 2017; Saudou and Humbert, 2016). Our results showed that HAP40 is a positive and essential regulator of HTT’s normal physiological functions in both flies and human cells (Fig. 4). Depletion of HAP40 could significantly compromise the neuroprotective activities of endogenous HTT. Lastly, our data suggest that HAP40-binding converts mutant HTT to a less toxic form (Fig. 8), as discussed above. This is probably due to the strong impact of HAP40 on the conformation and biochemical property of HTT protein. In particular, it is believed that for mutant HTT and other misfolding-prone proteins, their toxicity are closely associated with smaller aggregating species, while the large aggregates might be inactive or even protective (Arrasate et al., 2004; Ross and Poirier, 2005). Biochemically, it was shown that when expressed alone, full-length HTT exists in a conformational heterogeneous status, prone to oligomerize and aggregate (Huang et al., 2015), while HAP40-binding stabilizes HTT into a homogenous and monomeric globular structure (Guo et al., 2018). Thus, HAP40-binding converts HTT protein from aggregating-prone to a conformationally more stable and structurally more homogenous state. Therefore, HAP40 can directly affect the toxicity of mutant HTT by changing the course of its folding and aggregating dynamics, subsequently altering the composition and quantities of aggregating intermediates and the final aggregate species. In such a scenario, HAP40-free mHTT is toxic by itself, while HAP40-binding lowers it aggregating propensity and thereby converts it to a less toxic conformation. This double-edged roles of HAP40 on mutant HTT, reducing its potency of toxicity while increasing its total protein levels, might explain why despite several-fold increase of total HTT levels when co-expressed with HAP40, it did not translate into more severe neurodegeneration (Fig. 8E-I). Consistent with this hypothesis, the modifying effect by HAP40 is more specific for full-length HTT, not truncated HTT such as HTT exon 1 (Supplemental Fig. S3), likely due to the fact that HAP40 binds full-length but not truncated HTT (Guo et al., 2018; Pal et al., 2006).

Taken together, our results suggest that mutant HTT protein is toxic regardless of the presence of its binding partner HAP40, but when present at equal molarity, HAP40-free mutant HTT might be more potent in its toxicity than HAP40-bound mutant HTT. Therefore, HAP40 can exert profound and double-edged effect on HD pathogenesis by simultaneously affecting the normal physiological functions of HTT, the stability and overall levels of HTT, and the potency of mutant HTT toxicity *per se*. By selectively manipulating this multiplexity of HAP40 to preserve HTT’s neuroprotective activities while mitigate mutant HTT’s toxicity, this conserved and highly specific binding partner of HTT might potentially be an ideal candidate for novel therapeutic strategies such as “HTT-lowering” against HD.

## Figure Legends

**Supplemental Figure S1.**
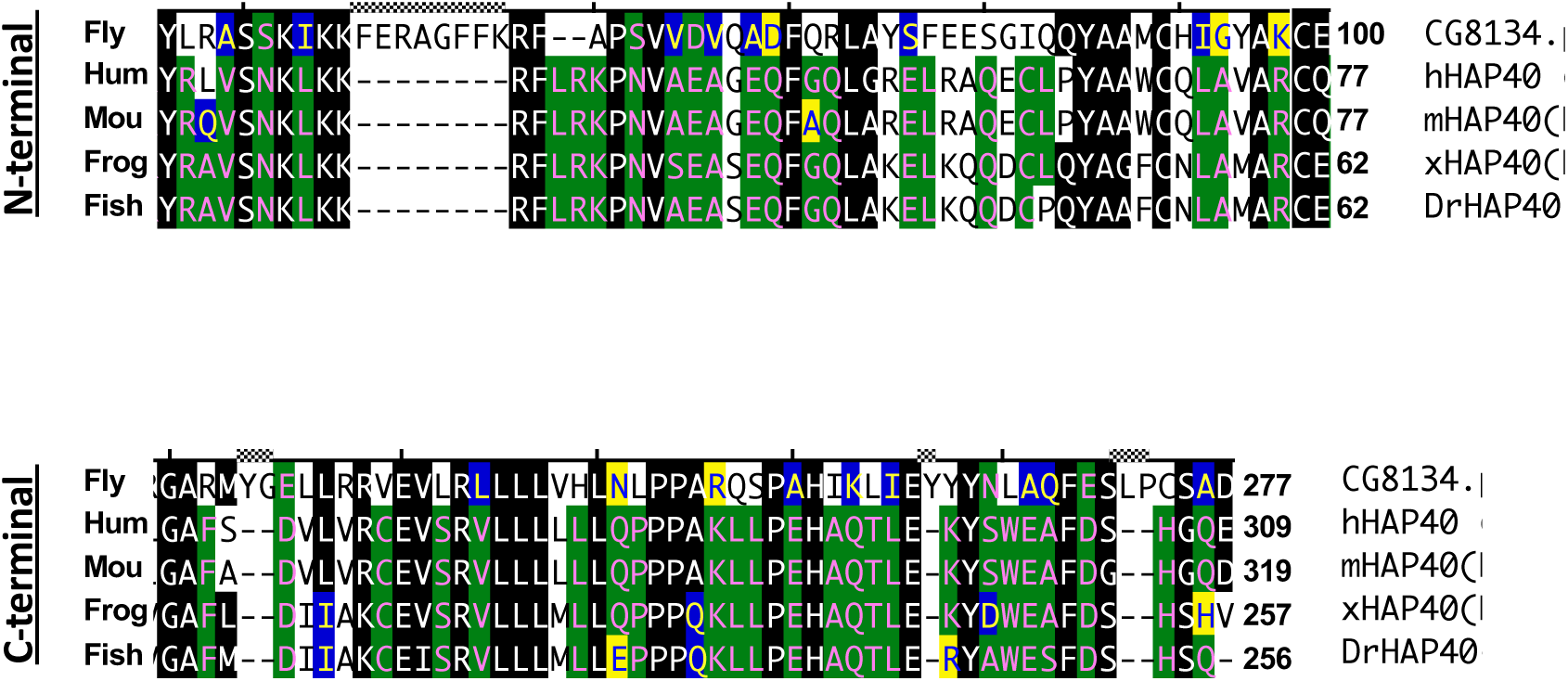
The N- and C-terminal regions of HAP40 are most conserved between CG8134 (dHap40) and vertebrate HAP40. Sequence alignment of the N- and C-terminal regions between CG8134 and HAP40 homologs from human, mouse, frog and zebra fish, as indicated. The amino acid positions of the corresponding boundaries are labeled accordingly. Amino acids that are identical to CG8134 are highlighted in black and with similar chemical properties highlight in color.

**Supplemental Figure S2.**
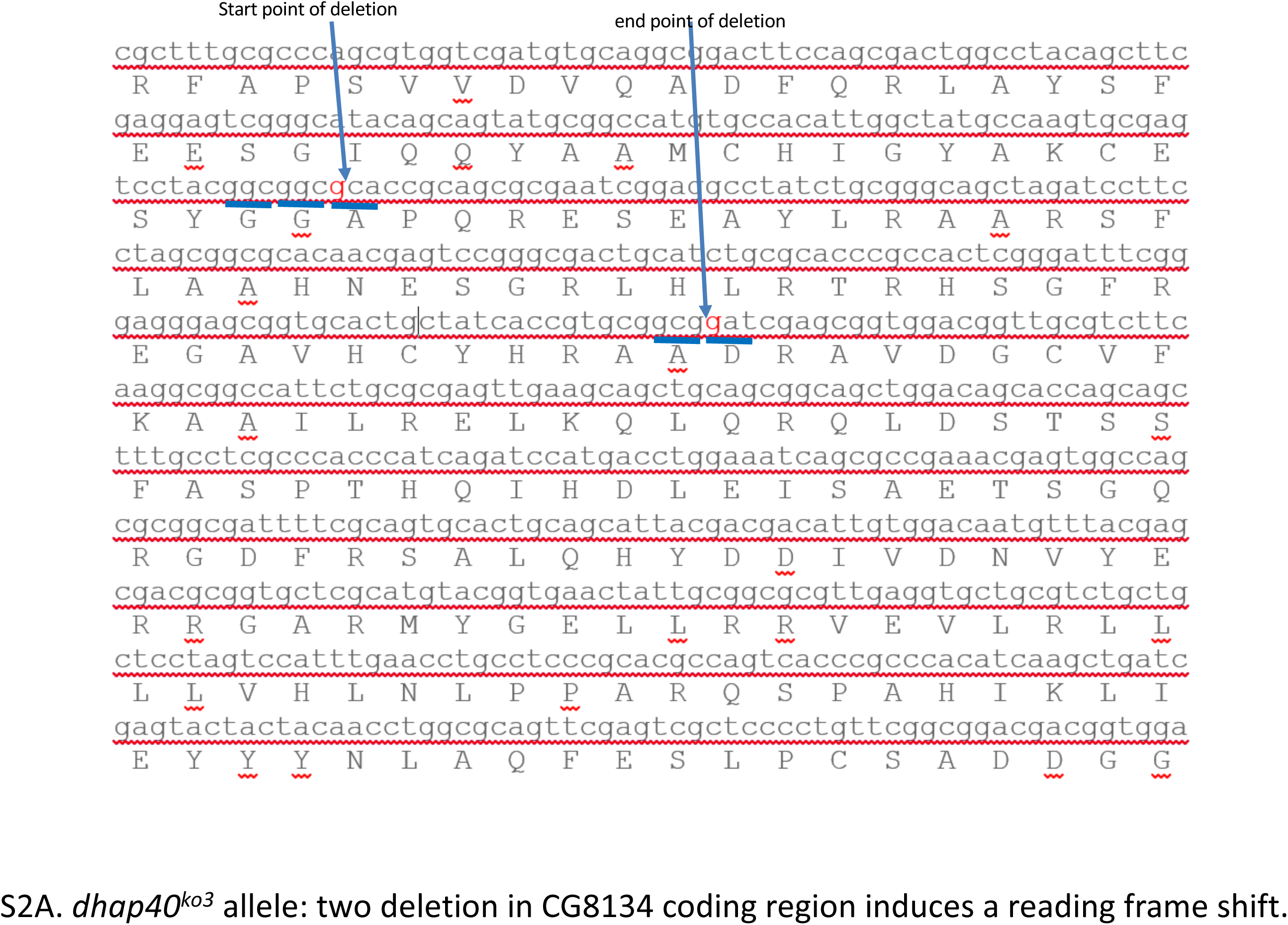
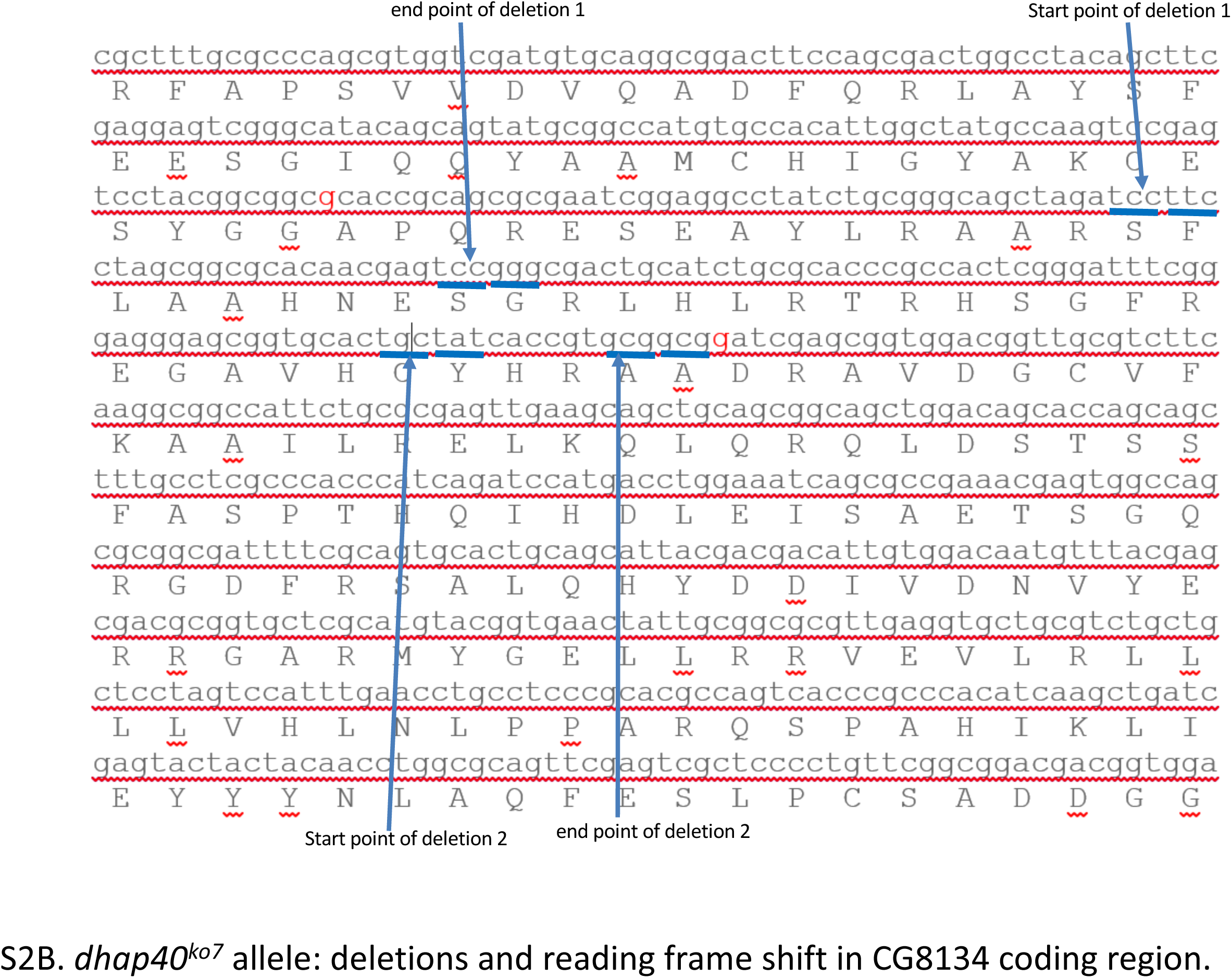
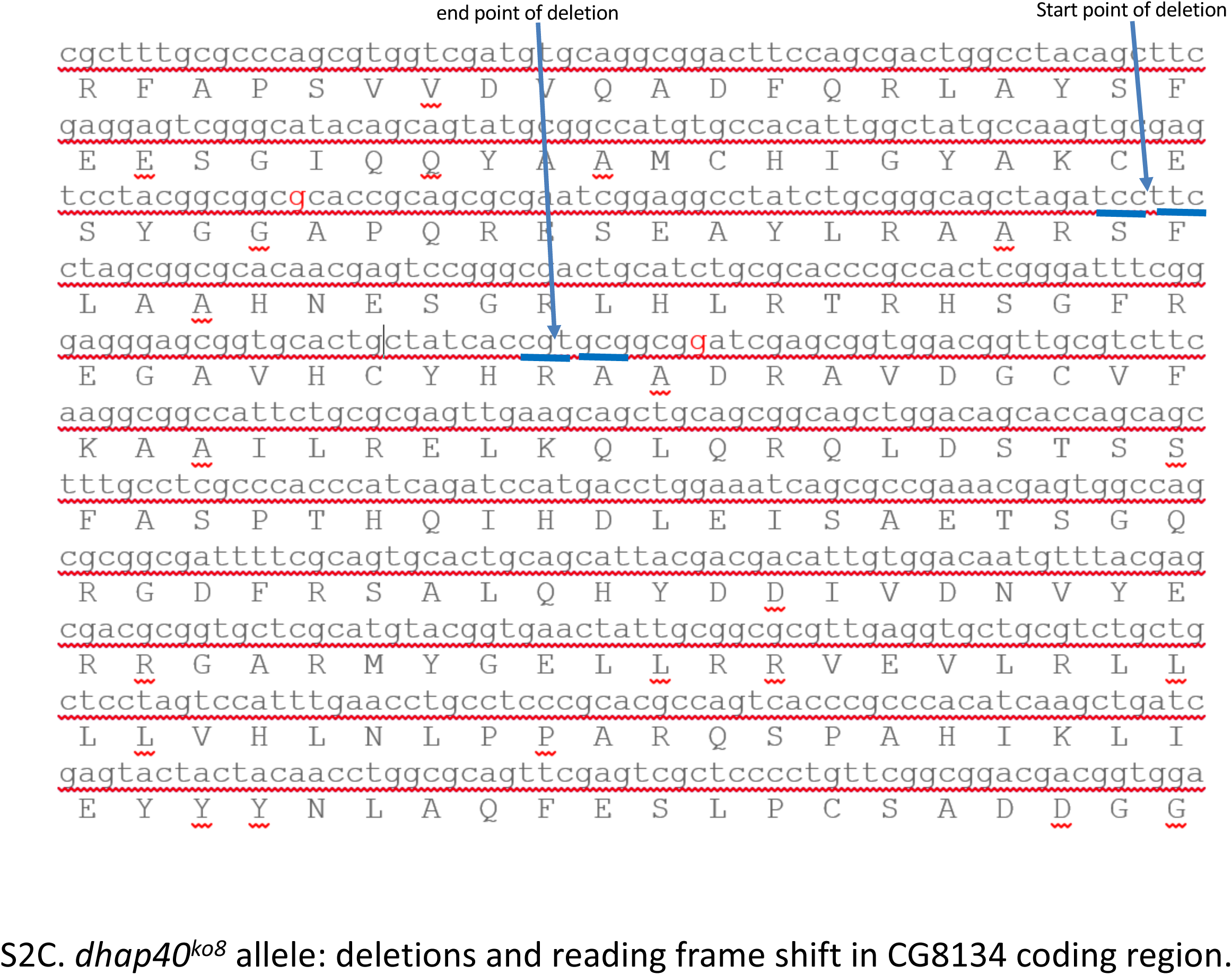
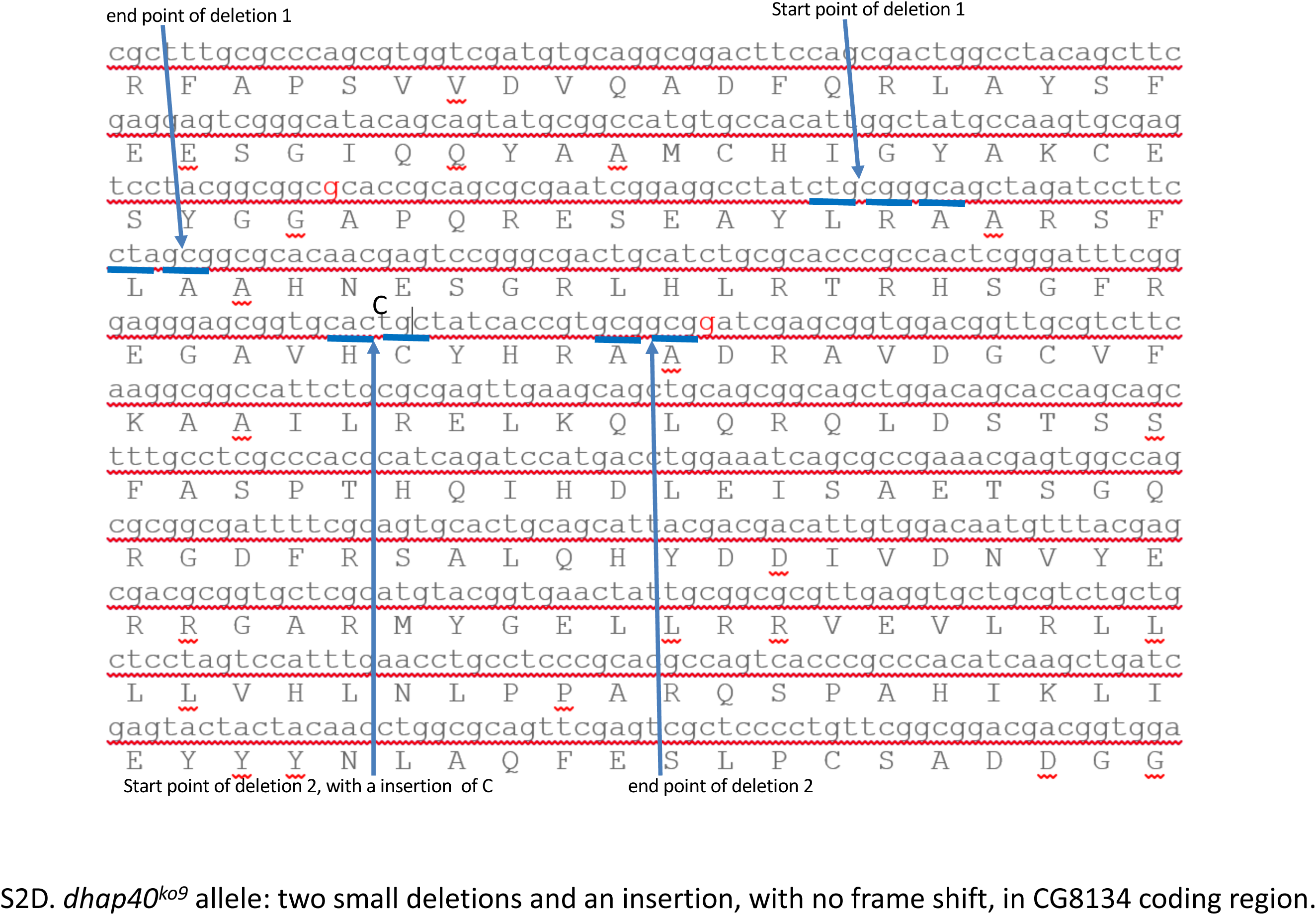
The molecular lesions in four established *dhap40* alleles. Genome sequence of exon 2 region of *cg8134* gene, in which the exact molecular lesions of the four validated *dhap40* mutant alleles, *ko3, ko7, ko8 and ko9*, are labeled in S2A-D, respectively, as indicated.

**Supplemental Figure S3.**
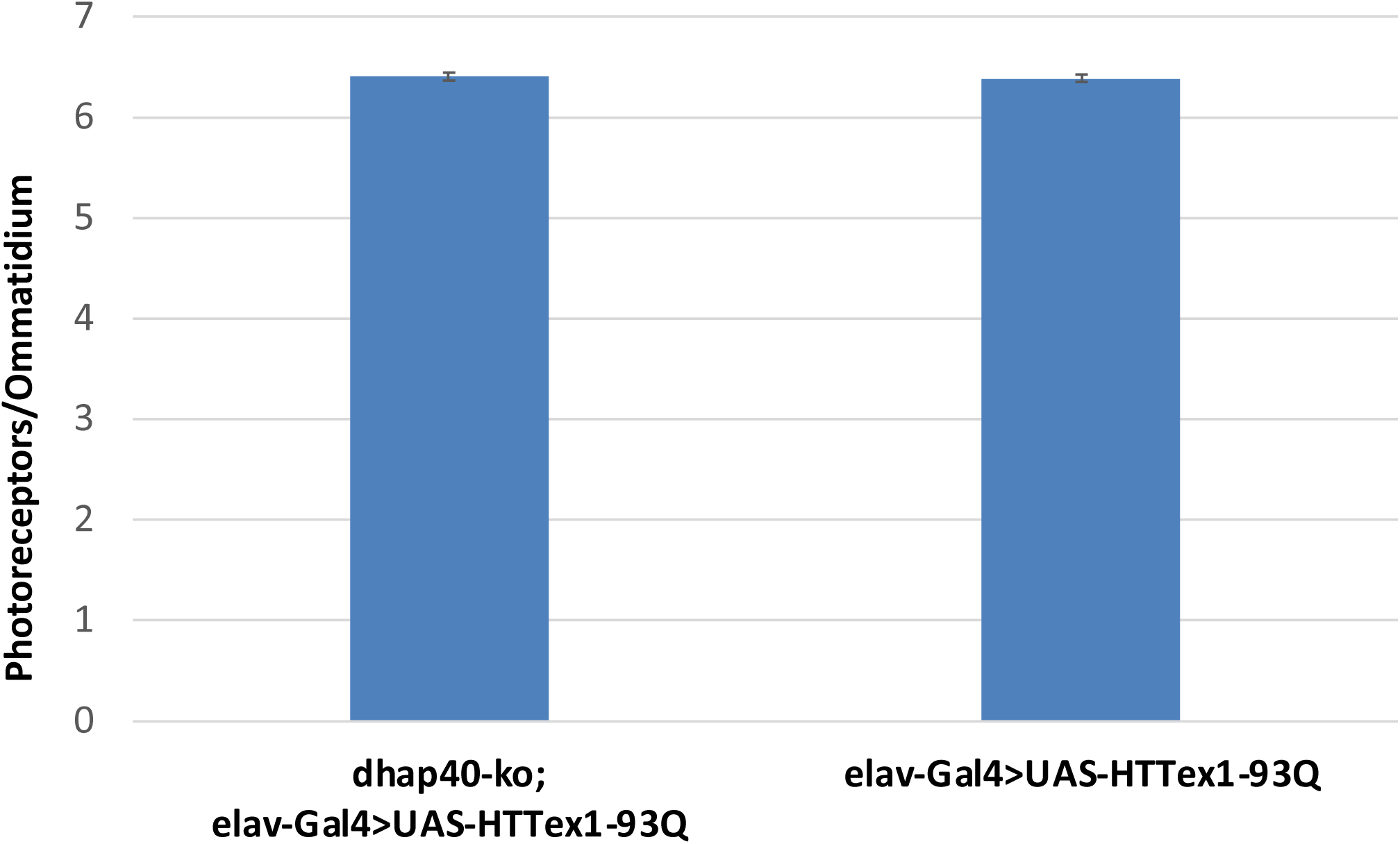
HAP40 does not modulate neurodegeneration induced by mutant HTT exon1 fragment in *Drosophila*. Loss of endogenous *dhap40* did not have apparent effect on neurodegenerative phenotypes induced by HTT-exon1-93Q. Bar chart presentation of the average number of intact photoreceptor cells (PRC) per ommatidium in 7-day-old female flies of the following genotypes: *“elav-*Gal4*/+; UAS-*HTT-exon1-93Q/+*”*: 6.4 PRC/ommatidium (n=12 flies); “*dhap40*^*ko3*^, *elav-*Gal4*/+; UAS-*HTT-exon1-93Q/+”: 6.4 PRC/ommatidium (n=12 flies). The difference between the two is insignificance (p=0.6 in rank-sum test). Flies in the study were cultured at 21°C.

## Materials and Methods

### *Drosophila* husbandry and genetics

Fly stocks were maintained at room temperature following standard culture conditions. All fly crosses were performed at 25°C following standard genetic procedure unless otherwise specified. The following fly lines were from Bloomington Drosophila Stock Center: UAS-Htt.128Q.FL (#33808), UAS-Htt.16Q.FL (#33810), UAS-cas9 (58985), elav-Gal4 (#458), daughterless-Gal4 (#55850), nSyb-Gal4 (#51635), nanos-Gal4 (#4442). UAS-Htt93Q-exon1 is a gift from Dr. Leslie Thompson.

### Genome tagging of *dhtt*

*dhtt* genome tagging constructs with C-terminal eGFP- and GA-TAP tags in pacman system were engineered following the established protocols (Venken et al., 2006; Venken et al., 2008). BacPac clone CH321-84B06 (BACPAC Resource Center (BPRC), Children’s Hospital Oakland Research Institute in Oakland, California), which covers all the genome coding regions of *dhtt* was selected as starting template for the tagging constructs (Venken et al., 2009). An 82Kb genome region covering *dhtt* gene, which contains all the coding region of dhtt in addition to 28.5kb upstream of ATG start codon and 15kb downstream of the stop codon of the encoded dHtt protein, was cloned by recombineering. DNA templates for eGFP and GS-TAP tags were amplified by PCR and used for recombineering essentially as described (Venken et al., 2009; Venken et al., 2008). The purified DNA for the corresponding pacman tagging constructs were injected into phiC31*integrase-expresssing* embryos with VK22 attP integrate site at second chromosome, and transgenes were selected and established following standard protocols. The expression of tagged transgenes were validated by Western blots on protein extracts from adult transgenic flies (Figures 1 and 2).

### Creating *dhap40* mutant alleles

The mutant *dhap40* lines were generated following the established procedures (Port and Bullock, 2016). Briefly, sgRNA1 (GGCAGCTAGATCCTTCCTAG) and sgRNA2 (GGTGCACTGCTATCACCGTG) targeting *cg8134* were cloned into pCFD6 vector and injected into fly embryo to generate corresponding transgenic lines. The resulting UAS-sgRNA 1, 2 were combined with UAS-cas9 and nanos-Gal4 for co-expression in fly germline. The male offsprings were selected and balanced to establish corresponding candidate mutant lines, and their genomic DNA were extracted and amplified using two primers: CAGCGTGGTCGATGTGCAG and CAGCGCCGAAACGAGTGG. For alleles with potential genome deletions, the protein extracts from the candidate homozygous flies were analyzed by Western blot assay to examine the absence of CG8134/dHap40 protein. The mutant lines with clear absence of CG8134/dHap40 protein were maintained for further characterization. Genome regions covering the established mutant alleles for *cg8134* gene were amplified by PCR followed by DNA sequencing to identify the molecular lesions within the *cg8134* gene. The confirmed *cg8134* null alleles were outcrossed with *w*^*1118*^ for 5 generations to establish stable fly lines with isogenic genetic background identical to the parental *w*^*11l8*^ line, before being employed for longevity and climbing assays.

### Molecular biology

pcDNA3-HTT-Q23-MYC (CH00038), pcDNA3-HTT-Q73-MYC (CH00039) and pcDNA3-HTT-Q145-MYC (CH00040) were from Coriell Institute deposited by the CHDI. The coding regions for HTT-Myc were released by select restriction enzymes and cloned into pUASTattB vector to generate corresponding transgenic fly lines.

For fly *cg8134*/*dhap40* overexpression, full-length cDNA clone for *cg8134* gene was digested from plasmid GH09650 (clone number 14278, Berkeley Drosophila Genome Project) by select restriction enzymes and cloned into pUAST vector to generate transgenic fly lines, or into pcDNA vector for transient expression by transfection in mammalian cells. For human HAP40 overexpression, full-length cDNA encoding human F8A1 (HAP40) was digested from pCS6(BC039693) plasmid (TCH1303, TransOMIC technologies) by restriction enzymes and cloned into pUAST vector for transgenic fly lines, or into pcDNA vector for transient expression by transfection in mammalian cells.

### Transgenic fly lines

For pUAST-HAP40 and pUAST-CG8134 (dHAP40) transgenes, the purified DNA were injected into *w*^*1118*^ embryos together with pπ25.7wc helper plasmid followed by standard transgenic procedures (Rubin and Spradling, 1982). For pUASTattB-HTT-Q23, Q73 and Q145 transgenes and pacman-dhtt-GS-TAP and pacman-dhtt-eGFP tagging constructs, the purified DNA constructs were injected into phiC31*integrase-expresssing* embryos with VK22 attP integrate site at second chromosome, followed by standard transgenic procedures (Bischof et al., 2007; Groth et al., 2004; Venken et al., 2006). Embryo injection and transgene selection were carried out by Genetivison Co. (Houston, Texas).

### Antibodies

Primary antibodies were from the following sources: mouse anti-actin (1:10000, MAB1501, Chemicon and #ab-6276, Abcam); mouse anti-Htt (MAB2166, Millipore. 1:1000); rabbit anti-F8A1/HAP40 antibody (HPA046960, Sigma), mouse anti-HA (12CA5, Roche); mouse anti-c-Myc (9E10, Santa Cruz); mouse anti-FLAG (M2, Sigma); rabbit anti-dHtt and rabbit anti-dHap40 (this study), rabbit anti-Atg8a and anti-Ref(2)p antibodies have been described previously (Zhu et al., 2017), chicken anti-GFP (Aves), mouse anti-SBP Tag (clone SB19-C4, Santa Cruz Biotechnology sc-101595).

Alexa680-(A-21076) and Alexa800-(926-32212) conjugated secondary antibodies for immunoblotting (1:10,000) were from Molecular probes-Invitrogen and LI-COR, respectively.

### Anti-dHtt and dHAP40 antibodies

To generate anti-dHtt antibody, the coding region corresponding amino acid 321-475 of dHtt protein was amplified by PCR and cloned into pET28b expression vector (Novagen). To generate anti-CG8134 antibody, the coding region of cg8134 gene was amplified by PCR and cloned into pET28b vector and verified by DNA sequencing. The constructs were transformed into BL21 E. Coli strain for inducible-expression of His-tagged proteins in bacteria. His-tagged dHtt and CG8134 protein fragments were purified by Nickle-beads following manufacturer’s instruction and used to immunize rabbits for antibody production (Covance). The specificities of the antibodies were verified in Western blot assays for the absence of the 400kDa dHtt, or 40kDa CG8134/dHap40 protein bands in whole protein extracts from the corresponding *dhtt-ko* and *dhap40-ko* mutant flies, respectively.

### Affinity purification of dHtt and associated proteins (dHaps) and MS-Proteomics

For protein extraction, 1 gram of overnight collections of fly eggs of the appropriate genotypes were mixed with 2ml of Lysis buffer (0.5% NP-40, 150mM KCl, 1mM EDTA 20mM Tris PH 7.5, with 1xProtease inhibitors and Phosphatase inhibitor cocktail (GenDepot), and homogenized by douncer. The total protein concentrations of the lysates were measured by Bradford method, and the lysates were dilute to a final protein concentration at about 15-20 mg/ml. After centrifuge at 100,000g (Beckman Optima TL Ultracentrifuge) for 20 minutes twice at 4°C to remove all debris from the lysate, the supernants were further processed according to the established procedures.

For GFP pulldown, 20 ul of slurry agarose GFP-Trap were added to the lysate, incubate for 60min at 4°C with constant rotation, followed by centrifuge to separate the supernant from the beads. The collected agarose beads were wash quickly two times use NETN buffer (0.5% NP-40, 170mM NaCl, 1mM EDTA, 50mM Tris PH 7.3), then eluate with 25ul 0.1M Glycine (PH 2.5). The elutes were to neutralize using 2.5ul 1M Tris (PH 7.0), followed by separation on 4-12% SDS-PAGE gradient gel (Nupage # 11062171-0033, Inviotrogen) and visualized by standard CB staining. The protein bands were excised and eluted for MS to identify the target proteins.

The GS-TAP purifications were essentially as described (Kyriakakis et al., 2008). Specifically, the cleared extract supernants were supplemented with 400ul of 50% rabbit IgG bead suspension (Sigma A2909), followed by incubation at 4°C for 2 hours with constant rotation. After binding, spin IgG beads down for 1 min at 500 g, wash the IgG beads with 10 ml of wash buffer (50 mM Tris pH 7.5, 5% glycerol, 0.2% IGEPAL, 1.5 mM MgCl_2_, 125 mM NaCl, 25 mM NaF, 1 mM Na_3_VO_4)_), then rinse with 400 ul TEV cleavage buffer (10 mM Tris pH 7.5, 100 mM NaCl,, 0.1% IGEPAL, 0.5 mM EDTA, 1 mM DTT), followed by TEV cleavage using 3 ul of AcTEV protease (Invitrogen, # 12575-015) in 160 ul of TEV cleavage buffer with constant rotation at 16°C for 90 minutes. After TEV cleavage, the elutes were separated from IgG beads, added to 120 ul of pre-washed Streptavidin beads (Pierce, #53117) and incubated at 4°C for 45 min with constant rotation, followed by washing with 6 ml 1x TEV cleavage buffer. After the last wash, elute with 200 ul of Elution buffer (2 mM biotin in 150mM NaCl, 10mM Tris Ph8.0, 0.5% NP-40).

### Cell culture and transient transfection

HEK293T or HeLa cells were maintained in DMEM medium (Mediatech) containing 10% fetal bovine serum (FBS, Invitrogen), 100 IU penicillin, and 100 µg/ml streptomycin. All cells were grown in a humidified incubator at 37°C with 5% CO2 and 95% air. Transfections of plasmid DNA were performed using Lipofectamine 2000 (Invitrogen) according to manufacturer’s instructions or for HEK293 cells by polyethylenimine (PEI) following standard procedure. For HTT and HAP40 protein degradation assays (Figure 6), cells were treated with 10 µM MG132, 10mM ammonium (NH4+) or 30uM chloroquine (CQ) for the duration of times as indicated. Cell lysates were prepared using NETN lysis buffer (20 mM Tris-HCl, pH 8.0, 100 mM NaCl, 0.5 mM EDTA, 0.5%(v/v) Nonidet P-40, 2.5 mM sodium pyrophosphate, 1 mM β-glycerolphosphate, 1 mM sodium orthovanadate, 1 µg/ml leupeptin, 1 mM phenylmethylsulfonyl fluoride) or CHAPS lysis buffer (CLB: 40 mM HEPES, pH 7.4, 2 mM EDTA, 10 mM pyrophosphate, 10 mM glycerophosphate, 0.3% CHAPS, protease inhibitors from Roche) as indicated.

### HTT and HAP40 knockout cell lines by CRISPR/Cas9

Single or double knockout cell lines for HTT and HAP40 in HEK293 and HeLa cells were created by CRISPR/Cas9 method essentially as described (Agudelo et al., 2017). For HTT-KO, the following sgRNAs were cloned into eSpCas9(1.1)_No_FLAG_ATP1A1_G2_Dual_sgRNA vector (Addgene): HTT-sgRNA1 (GAAGGACTTGAGGGACTCGA) and HTT-sgRNA4 (ATGACGCAGAGTCAGATGTC), or HAP40-SgRNA (GTGGCCAGGAGCCTCCGCCC). The corresponding constructs were transfected into HEK293 or HeLa cells, followed by Ouabain selection three days after transfection and expansion of single cell clones in 24-well plates as described (Agudelo et al., 2017). The established KO cell lines were validated by Western blot assays with anti-HTT (mAb2166, Millipore) or anti-HAP40 (HPA046960, Sigma) antibodies for the absence of endogenous HTT or HAP40 proteins, respectively.

### Bright field imaging of adult fly eyes

Adult flies of appropriate genotypes were orientated on glass slides with nail polish. Z-stack scanning of adult eyes were recorded under the 10X objective using Zeiss Axioimager Z1 microscope (usually 10-20 layers scanned) and reconstructed into 3D projection using CZFocus software.

### Climbing assay

Climbing assays were performed as described previously (Zhang et al., 2009). Briefly, 20 female flies were placed into a clear fly vial, gently banged to the bottom, and then given 18 seconds to climb a 5 cm vertical distance. Flies that successfully do so were counted and their ratio at each time point were analyzed by t-test. Each fly group was tested every 7 days and transferred into a new vial with fresh food every 3-4 days.

### Longevity assay

Viability assays were performed as described previously (Zhang et al., 2009). Briefly, 20 newly hatched male or female flies of a specific genotype were placed into individual vials with fresh fly food at 25°C. The number of dead flies was recorded every 3 days until all flies died. Flies were transferred into a new vial with fresh food every 3 days to prevent them from stuck to old food or become dehydrated. The survival curve for each genotype were analyzed by log-rank test.

### Pseudopupil assay of adult fly eyes

Pseudopupil assay of adult fly eyes were performed as described (Franceschini, 1972). Briefly, flies of a specific genotype were oriented and affixed in nail polish. The eyes were immersed in lens oil (Resolve Microscope Immersion oil, Thermo Scientific, Cat# M5000) and imaged through a 40X oil microscope lens (Zeiss, AX10). For each fly head, the rhabdomeres in 15-20 ommatidia were counted. Usually, 10-20 flies for each genotype were analyzed by Mann-Whitney rank sum test.

### Biochemistry

#### Western blotting

Standard 10% to 14% SDS-PAGE gels were used for separation of most proteins except for HTT, which was better analyzed by NuPAGE Tris-Acetate gels from Invitrogen specially formulated for detection of proteins with large molecular weight. The boiled samples were separated on SDS-PAGE and transferred to nitrocellulose membranes from Millipore. After blocking with 5% nonfat milk in Tris-buffered saline with 0.1% Tween-20 for 1 hour, membranes were incubated with primary antibodies. Secondary antibodies conjugated with Alexa-800 or Alexa-680 (Invitrogen) or HRP (KPL) were used and the signals were detected by the Odyssey Infrared Imaging System and quantified by Odyssey Application Software 3.0 or by densitometry of the digital images using ImageJ software (NIH).

#### Co-immunoprecipitation (co-IP)

Co-IP were performed as described. Specifically, transiently transfected HEK293T cells in 100 mm dishes were lysed in a NETN or CLB as indicated, sonicated three times for 5 sec each, and centrifuged at 14,000 rpm for 30 min at 4°C. Epitope-tagged proteins or endogenous proteins were immunoprecipitated from the cell lysate with anti-HA, anti-Myc, anti-FLAG, anti-Htt, and anti-HAP40 antibodies and Protein A/G Plus-agarose beads (sc-2003, Santa Cruz) as indicated. Immunoprecipitates or whole cell extracts (WCE) were analyzed by standard Western blotting.

#### Autophagic measurements

All autophagy assays were carried out as previously described (Bjorkoy et al., 2009; Kimura et al., 2009; Klionsky et al., 2016). For LC3 lipidation assay, cells were harvested in 2% Triton X-100/PBS buffer containing protease inhibitors for maximum LC3-II extraction according to the previous studies (Kimura et al., 2009). LC3-II and loading control actin were detected by rabbit anti-LC3 antibody (MBL international) and mouse anti-actin antibody (Chemicon), respectively. The level of LC3 lipidation was quantified as the ratio of the measured LC3-II to actin levels.

### Statistics analysis and data acquisition

The statistical significance of the difference between experimental groups was determined by two-tailed unpaired Student’s *t*-test, or one-way analysis of variance followed by Bonferroni *post hoc* test. Where multiple comparisons were performed, we used normalization to control values. Differences were considered significant for *P*<0.05 (noted in the figures as *). Data are presented as mean+s.e.m from a minimum of three independent experiments. The exact sample size (n) is indicated in each figure and it corresponds to individual experiments unless otherwise stated. All the experiments were done at least 3 times and in duplicate or triplicate to account for technical variability.

For the studies in *Drosophila*, sample group allocation was based on genotype and the genotypes were blinded to the observed except for those cases in which tissues from different animals have to be pooled.

## Acknowledgements

We are grateful to Dr. Juan Botas for UAS-hHtt fly lines, Dr. Alexey Veraksa for pUAST-TAP tagging constructs, Dr. Hugo Bellen for *dhtt* genome DNA clones and technical support, Matthew Zamarripa from Rice University for blind test to count ommatidia in adult fly eyes, and Bloomington Drosophila Stock Center for *Drosophila* lines. This work was supported by NIH grant R01 NS110943 (to S.Z.).

## Author contributions

SY Xu performed most of the studies on *cg8134/dhap40* gene in *Drosophila*, including generating and analyzing transgenic flies for F8A1/HAP40, cg8134/dhap40 and HTT-Q23, Q73 and Q145, dHap40 antibody characterization, creation of *dhap40* mutant lines by CRISPR and their phenotypic characterizations in normal and fly HD models. G Li performed most of the studies on HAP40 in mammalian cells, including generated HTT-KO and HAP40-KO cell lines, co-IP studies, autophagy assays, protein expression and turnover assays for HAP40 and HTT. DS Chen and ZH Chen performed affinity purification studies for dHtt and associated proteins from fly embryos carrying *dhtt-GFP* and *dhtt-GS-TAP* genome transgenes; Z Xu generated the GS-TAP and GFP-tag genome tagging constructs for *dhtt* gene; X Ye analyzed the expression of HTT and HAP40 proteins in extracts of whole fly tissues of different genetic background. L Ye contributed to *Drosophila* genetic studies; EF Stimming and D Marchionini contributed to experiment design and result discussion. S Zhang designed experiments, coordinated the study, analyzed data and wrote the manuscript.

